# A whole transcriptomic approach reveals novel mechanisms of organophosphate and pyrethroid resistance in *Anopheles arabiensis* from Ethiopia

**DOI:** 10.1101/2021.07.09.451871

**Authors:** Louisa A. Messenger, Lucy Mackenzie Impoinvil, Dieunel Derilus, Delenasaw Yewhalaw, Seth Irish, Audrey Lenhart

## Abstract

The development of insecticide resistance in malaria vectors is of increasing concern in Ethiopia because of its potential implications for vector control failure. To better elucidate the specificity of resistance mechanisms and to facilitate the design of control strategies that minimize the likelihood of selecting for cross-resistance, a whole transcriptomic approach was used to explore gene expression patterns in a multi-insecticide resistant population of *Anopheles arabiensis* from Oromia Region, Ethiopia. This field population was resistant to the diagnostic doses of malathion (average mortality of 71.9%) and permethrin (77.4%), with pools of survivors and unexposed individuals analyzed using Illumina RNA-sequencing, alongside insecticide susceptible reference strains. This population also demonstrated deltamethrin resistance but complete susceptibility to alpha-cypermethrin, bendiocarb and propoxur, providing a phenotypic basis for detecting insecticide-specific resistance mechanisms. Transcriptomic data revealed overexpression of genes including cytochrome P450s, glutathione-s-transferases and carboxylesterases (including CYP4C36, CYP6AA1, CYP6M2, CYP6M3, CYP6P4, CYP9K1, CYP9L1, GSTD3, GSTE2, GSTE3, GSTE4, GSTE5, GSTE7 and two carboxylesterases) that were shared between malathion and permethrin survivors. We also identified nineteen highly overexpressed cuticular-associated proteins (including CYP4G16, CYP4G17 and chitinase) and eighteen salivary gland proteins (including D7r4 short form salivary protein), which may be contributing to a non-specific resistance phenotype by either enhancing the cuticular barrier or promoting binding and sequestration of insecticides, respectively. These findings provide novel insights into the molecular basis of insecticide resistance in this lesser well-characterized major malaria vector species.

**Importance:** Insecticide-resistant mosquito populations remain a significant challenge to global malaria vector control. While substantial progress has been made unraveling resistance mechanisms in major vector species, such as *Anopheles gambiae* and *An. funestus*, comparatively less is known about *An. arabiensis* populations. Using a whole transcriptomic approach, we investigated genes associated with resistance to insecticides used to control *An. arabiensis* in Ethiopia. Study findings revealed shared detoxification genes between organophosphate- and pyrethroid-resistant vectors and highly overexpressed cuticular-associated proteins and salivary gland proteins, which may play a role in enhancing insecticide resistance. The whole transcriptomic analysis detected novel resistance-associated genes, which warrant functional validation to determine their specificity to particular insecticides and their potential to confer cross-resistance between different insecticides with the same mode of action. These genes may contribute to the development of diagnostic markers to monitor insecticide resistance dynamics in the field.

## Introduction

Globally, malaria mortality has fallen since 2010, largely due to the scale-up of diagnosis, treatment and insecticide-based vector control interventions. However, since 2016, the rates of decline have stalled in the World Health Organization regions of Africa, Southeast Asia and the Western Pacific and even reversed in the Eastern Mediterranean and the Americas (1). Concurrently, insecticide resistance among major malaria vector species has become widespread, affecting approximately 90% of countries with ongoing malaria transmission (1) and threatening vector control efforts worldwide.

In Ethiopia, insecticide resistance in the principal malaria vector species *Anopheles arabiensis* has been a public health concern for decades. Indoor residual spraying (IRS) using DDT was first implemented in 1959, and insecticide-treated net (ITN) distribution was initiated in 1997 and scaled up since 2005 (2). Following the detection of DDT resistance in 2009, DDT was replaced with deltamethrin for IRS, initially alongside bendiocarb from 2011 until 2013, after which bendiocarb and propoxur were sprayed in different geographical areas. In 2015, pirimiphos-methyl was introduced and is now used alongside propoxur across the country (3). In parallel, more than 80 million pyrethroid-treated long-lasting insecticidal nets (LLINs) have been distributed in Ethiopia since 2008 (2). This heterogeneous use of different chemicals has resulted in highly focal, dynamic resistance patterns across Ethiopia, broadly reflecting longitudinal shifts in the national insecticide policy (3,4). Populations of *An. arabiensis* are now largely resistant to DDT and deltamethrin, with reduced susceptibility to malathion, pirimiphos-methyl, propoxur and bendiocarb reported in some locations (7,8). The presence of the L1014F-*kdr* allele was first reported from areas surrounding the Gilgel-Gibe hydroelectric dam in southwestern Ethiopia in 2010 (5). In these populations, L1014F-*kdr* was practically fixed and this target site mutation is now commonly detected elsewhere in Ethiopia at varying frequencies (3). Elevated levels of glutathione-S-transferases (GSTs) have also been observed in some *An. arabiensis* populations from Oromia and Benishangul-Gumuz regions (8). To date, other target site mutations, including L1014S-*kdr*, N1575Y and G119S-*Ace-1*, have not been detected in Ethiopia (3,4).

In Oromia region, *An. arabiensis* has demonstrated resistance to insecticides belonging to four of the chemical classes historically used for adult vector control (pyrethroids, carbamates, organophosphates and organochlorines) (3,4). In this area, restoration of susceptibility following pre-exposure to the synergist piperonyl butoxide (PBO) (3,6), coupled with a lack of association between phenotypic resistance and L1014F-*kdr* frequency and the complete absence of other target-site mutations (L1014S-*kdr*, N1575Y and G119S-*Ace-1*), suggest that metabolic mechanisms may play an important role in resistance (3,4).

In African *Anopheles*, several cytochrome P450 monooxygenases (CYP450s), carboxylesterases (COEs) and GSTs, have been functionally associated with pyrethroid resistance (7-11). In addition to detoxification enzymes, other gene families, including α-crystallins, hexamerins and ATP synthases (12), Maf-S, Dm and Met transcription factors (12,13), D7r2 and D7r4 salivary gland proteins (14), a sensory appendage protein, SAP2 (15) and cuticular proteins (16) have been associated with insecticide resistance. While over-expression of a number of these proteins is conserved across countries and sub-species of the *An. gambiae* s.l. complex (12), there is still a considerable paucity of data regarding mechanisms of resistance in *An. arabiensis*, especially in Ethiopia (3,4,17). Currently, only CYP6P4 and GSTD3 have been directly linked to local deltamethrin and DDT resistance (17).

In Ethiopia, nationwide insecticide resistance management strategies rely on the tactical deployment of IRS and LLINs with differing active ingredients. For such strategies to succeed, there needs to be a clear understanding of the specificity of resistance mechanisms to individual insecticides and the likelihood of selecting for cross-resistance mechanisms. To improve our understanding of these factors in *An. arabiensis*, we undertook a whole transcriptomic approach to characterize gene expression patterns in a multi-insecticide resistant field population of *An. arabiensis* from south-west Ethiopia.

## Results

### Phenotypic insecticide resistance

Indoor resting F_0_ adult *An. gambiae* s.l. were collected from houses in Asendabo, Oromia region, Ethiopia from July-September 2017 and F_1_ progeny were generated by forced-oviposition (18). Susceptibility to the diagnostic doses (1X) of malathion (organophosphate) and permethrin (pyrethroid) was determined for 273 F_1_ *An. gambiae* s.l. mosquitoes, using U.S. Centers for Disease Control and Prevention (CDC) bottle bioassays (20). These mosquitoes were subsequently confirmed via species-specific PCR as *An. arabiensis* (19). The average mortality to malathion was 71.9% [95% CI: 65.3-78.5] and to permethrin was 77.4% [95% CI: 44.0-100.0%]. Resistance intensity assays, using an additional 1183 PCR-confirmed F_1_ *An. arabiensis*, were conducted with alpha-cypermethrin (1X), bendiocarb (1X), propoxur (1X), deltamethrin (1X, 2X, 5X and 10X) and permethrin (1X, 2X, 5X and 10X) (20). Complete (100%) mortality was observed to the diagnostic doses of alpha-cypermethrin, bendiocarb and propoxur, while moderate to intense resistance was detected to deltamethrin and permethrin, with small proportions of mosquitos capable of surviving five to ten times the diagnostic concentrations (Figure 1).

**Figure 1.**
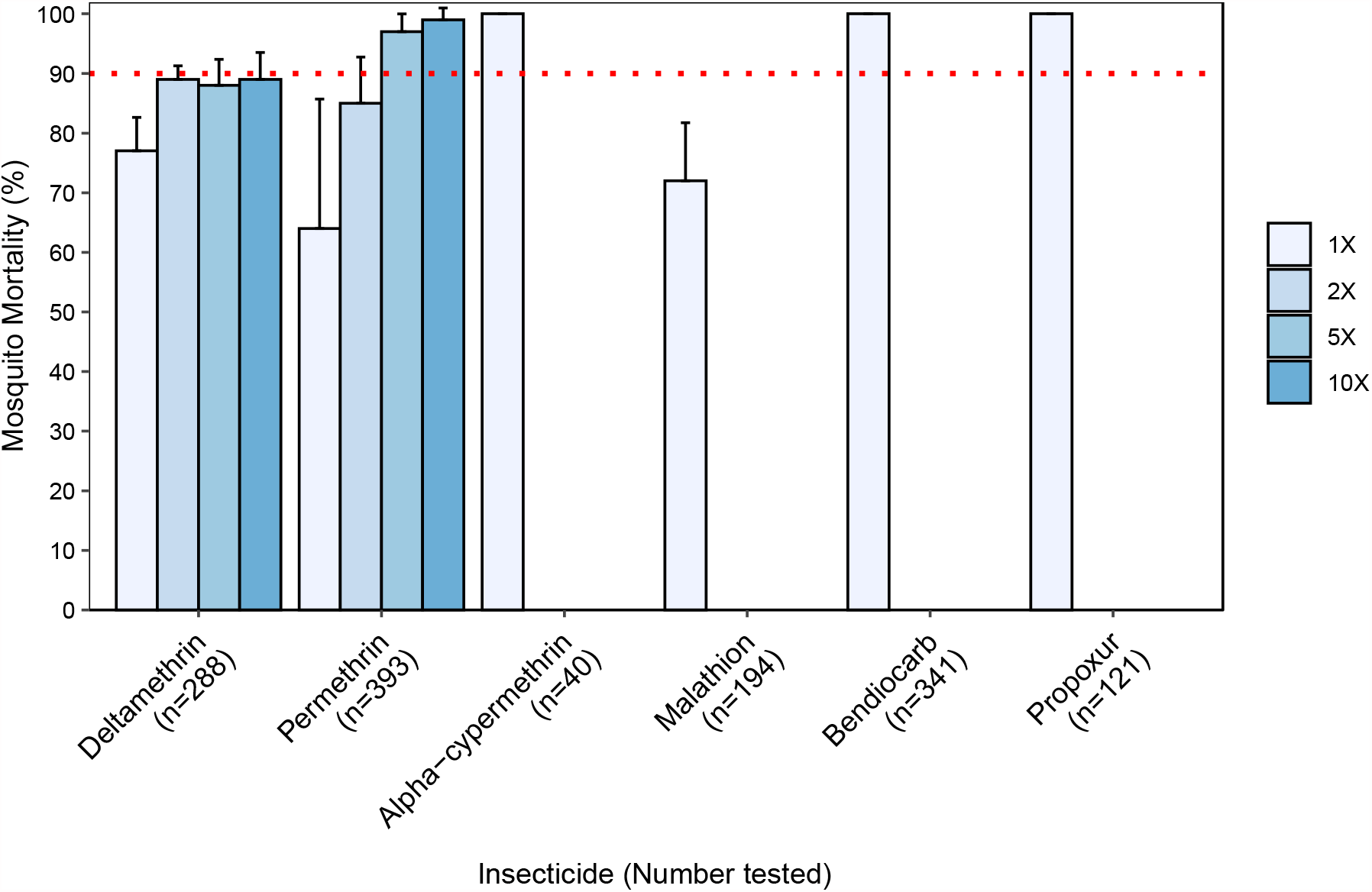
Bottle bioassay results for pyrethroid (deltamethrin, permethrin and alpha-cypermethrin), organophosphate (malathion) and carbamate (bendiocarb and propoxur) insecticides among *An. arabiensis* from Asendabo, Ethiopia. Bars show the mean mortality after 30 minutes of insecticide exposure across bottle replicates with 95% confidence intervals. The red dashed line indicates the threshold of 90% mortality, below which a population is considered resistant.

### Target site mutations

Phenotyped individuals were screened for known insecticide resistance target site mutations. The G119S-*Ace-1* mutation was not detected in any mosquitoes from the malathion bioassays (n=173). The L1014F-*kdr* mutation was identified in 52% (30/58) of *An. arabiensis* exposed to the diagnostic dose of permethrin, with allele frequencies of 0.65 in surviving mosquitoes and 0.26 in dead mosquitoes. A greater proportion of *An. arabiensis* surviving permethrin bioassays were homozygous for L1014F-*kdr* (46%; 6/13) compared to those that died (9%; 4/45), and 38.5% of survivors (5/13) and 33% of dead individuals (15/45) were heterozygous. No deviations from Hardy-Weinberg equilibrium were observed in either surviving or dead mosquitoes (χ^2^=0.29; *p*=0.59 and χ^2^=0.69; *p*=0.41, respectively). The L1014S-*kdr* allele was not detected in any sample tested.

### RNA sequencing quality control and mapping metrics

Malathion or permethrin bioassay survivors, field mosquitoes which were not exposed to insecticide, and two *An. arabiensis* susceptible reference strains (originally from Sudan or Ethiopia – Dongola or Sekoru, respectively) were submitted for transcriptomic analysis. For the malathion experiment, Illumina RNA-sequencing generated more than 620 million raw reads across three biological replicates, sequenced in technical duplicate with an average of 68.9 (± 5.1) million reads per group. (Table S1). After filtering and quality trimming, an average of 67.6 (± 5.0) million reads were retained per group (98.15%) for subsequent analysis. An average of 51 (± 7.8) million quality filtered reads per group (75.40%) were mapped to the whole *An. arabiensis* Dongola AaraD1.11 reference genome, with around 59% of the counted fragments mapped to all exonic features of the gene set (Table S1). The permethrin experiment generated more than 569 million reads across three biological replicates, sequenced in technical duplicate with an average of 63.3 (±10.9) million reads per group (Table S1). Quality control filtering retained an average of 61.4 (±10.7) million reads per population (97.02%), with an average of 42.6 (±14.3) million total filtered reads aligned to the reference genome (69.48%) and around 64 % of the counted fragments successfully assigned to exons of the gene set (Table S1). Full results for the analyses of the malathion and permethrin experiments are presented in Table S2, and results of gene ontology (GO) enrichment analysis for sets of differentially expressed genes (DEGs) are shown in Table S3.

### Differentially expressed genes associated with malathion resistance

Differential expression analysis was performed on transcripts retained after quality control and removal of genes with low read counts. Aligned reads were mapped to the *An. arabiensis* genes dataset (AaraD1.11) to quantify levels of gene expression, with between 52-69% of alignments successfully assigned to the exonic regions of the reference genome (Table S4). Three pairwise comparisons were conducted for malathion: resistant *vs* susceptible (R-S; MAL-R *vs* DON), resistant *vs* unexposed control (R-C; MAL-R *vs* CON-M) and unexposed control *vs* susceptible (C-S; CON-M *vs* DON). The R-C comparison allowed us to account for induction of transcription during insecticide exposure; genes were filtered by analysing their expression profiles in the susceptible Dongola strain, with the assumption that constitutive resistance genes will be significantly differentially expressed between both bioassay survivors and the non-exposed field mosquitoes, when compared to the susceptible strain.

At the most conservative level (*P*-values adjusted for multiple testing based on a false discovery rate (FDR)<0.01 and fold change (FC)>2), a total of 1212 (12.2%; 872 upregulated and 340 downregulated) genes were significantly differentially expressed in mosquitoes that survived malathion exposure and 598 (6.0%; 398 upregulated and 200 downregulated) were significantly differentially expressed in non-insecticide exposed field mosquitoes as compared to the susceptible strain (Figure 2A; Table 1). A total of 170 (1.8%; 137 upregulated and 33 downregulated) genes were significantly differentially expressed in mosquitoes that survived malathion exposure compared to their non-insecticide exposed counterparts (Figure 2A; Table 1).

**Table 1.** Summary of differential gene expression analyses for malathion and permethrin experiments.

| Condition | # of genes tested | DE genes (adjP<0.05) |  | DE genes (adjP<0.01) |  | DE genes(\|FC\|>2 & adjP<0.05) |  | DE genes(\|FC\|>2 & adjP<0.01) |  |
| --- | --- | --- | --- | --- | --- | --- | --- | --- | --- |
|  |  | Up | Down | Up | Down | Up | Down | Up | Down |
| MAL-R *vs* CON-M | 9609 | 455 | 163 | 209 | 59 | 203 | 61 | 137 | 33 |
| MAL-R *vs* DON | 9959 | 1998 | 1586 | 1557 | 1027 | 893 | 364 | 872 | 340 |
| CON-M *vs* DON | 9906 | 661 | 972 | 392 | 616 | 229 | 456 | 398 | 200 |
| PERM-R *vs* CON-P | 9669 | 595 | 424 | 351 | 246 | 214 | 192 | 179 | 155 |
| PERM-R *vs* SEK | 9293 | 2471 | 2640 | 1955 | 2005 | 1083 | 1156 | 1057 | 1126 |
| CON-P *vs* SEK | 9752 | 2790 | 2745 | 2259 | 2259 | 1193 | 2790 | 1153 | 1159 |
| MAL-R *vs* PERM-R | 9551 | 139 | 173 | 68 | 94 | 60 | 102 | 45 | 77 |
| DON *vs* SEK | 9961 | 3074 | 3148 | 2579 | 2619 | 1564 | 1447 | 1557 | 1414 |
| PERM-R *vs* DON | 9885 | 1565 | 1256 | 1086 | 754 | 714 | 450 | 673 | 401 |
| CON-P *vs* DON | 9999 | 1354 | 1046 | 981 | 645 | 632 | 343 | 594 | 295 |

DON=Dongola susceptible colony; MAL-R=alive after malathion exposure; PERM-R=alive after permethrin exposure; SEK=Sekoru susceptible colony; CON-M=field population not exposed to insecticide in malathion experiment. CON-P=field population not exposed to insecticide in permethrin experiment.
DE=differentially expressed; FC=fold change; adjP=*P*-value adjusted for multiple testing (22).

**Figure 2.**
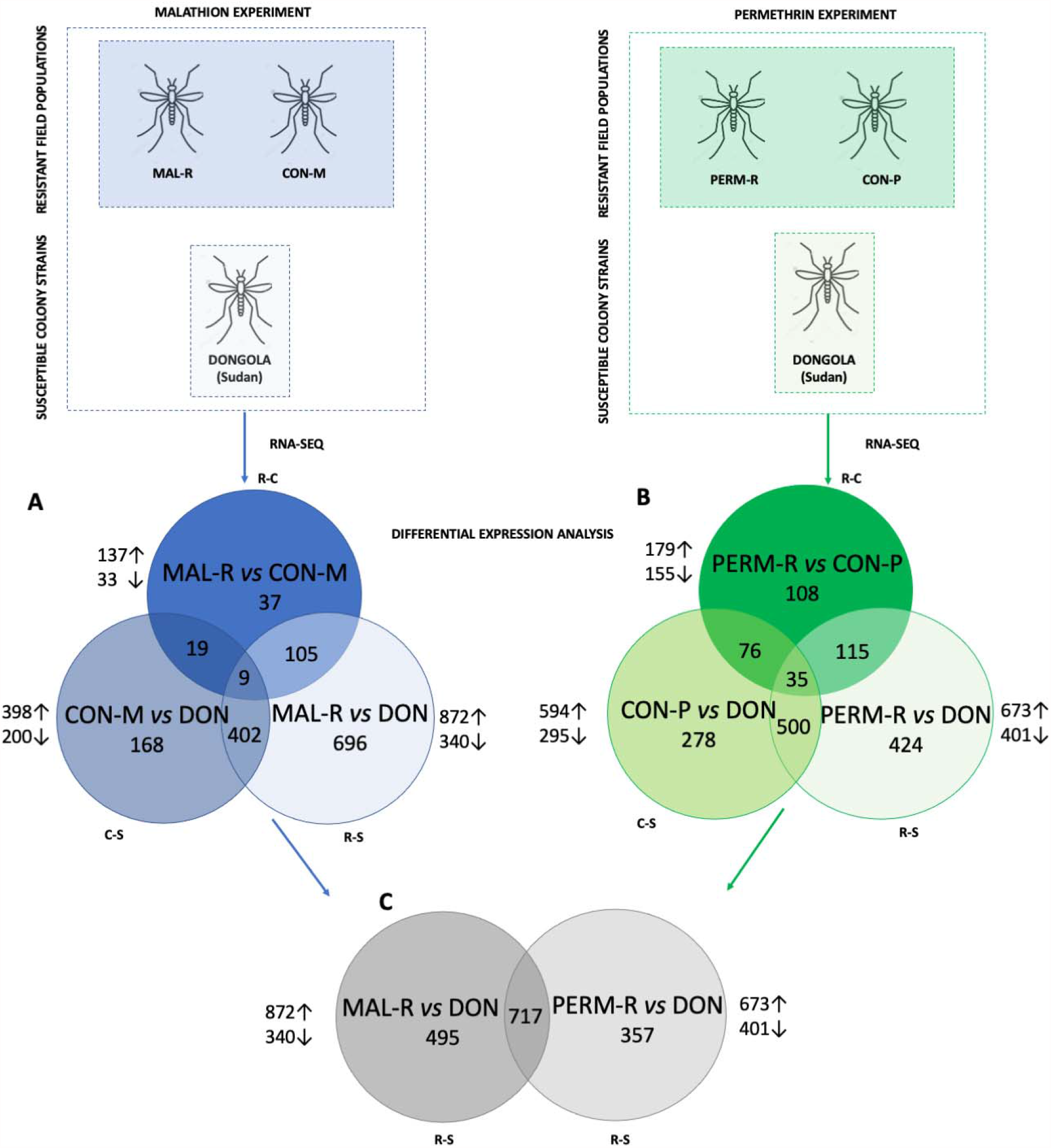
Experimental design and differentially expressed genes among resistant (R), susceptible (S) and unexposed (C) mosquito populations in malathion (A) and permethrin (B) experiments and in both (C). Each Venn diagram section shows the number of differentially expressed genes meeting each set of conditions (*P*-values were adjusted for multiple testing based on FDR<0.01 and FC>2). For a list of all DEGs for each comparison see Table S2.

Of the genes that were differentially expressed in all treatment groups (n=9), 2 were upregulated while 7 were downregulated in one or more conditions (Figure 2A). Five of these genes had retrievable annotations, all of which were molecular functions or cellular components (for R-C/R-S/C-S comparisons: AARA017080=peptide methionine sulfoxide reductase, FCs=2.57, 0.43 and 0.18; AARA016556=sulfotransferase, FCs=2.23, 23.88 and 9.92; AARA007045=protease M1 zinc metalloprotease, FCs=0.40, 0.18 and 0.44; AARA002630=transient receptor potential protein, FCs=0.21, 0.49 and 2.37; and AARA002503=ion binding protein, FCs=0.37, 0.04 and 0.17, respectively).

A total of 405 genes were differentially expressed commonly in the R-S and C-S groups (Figure 2A). Among the top 10 over-expressed genes with retrievable annotations were enzymes with structural, cellular or immune functions, including chitinase (AARA007329: FCs=50.04 and 10.80 for R-S/C-S comparisons, respectively), D7r4 short form salivary protein (AARA016237: FCs=33.29 and 31.34), cytoplasmic actin (AARA015772: FC=29.53 and 7.33), cuticular protein CPLCG (AARA011115: FCs=26.80 and 20.12), alkaline phosphatase (AARA002132: FCs=26.33 and 11.83), sulfotransferase (AARA016556: FCs=23.88 and 9.92), serine protease (AARA009441: FCs=23.73 and 24.43), polyubiquitin (AARA016579: FCs=21.67 and 31.07), ADP/ATP carrier protein (AARA017958: FCs=21.15 and 5.23) and deoxyribonuclease (AARA000505: FCs=17.0 and 12.15). A total of 19 genes were differentially expressed commonly in the R-C and C-S groups (Figure 2A). Among the top over-expressed genes with retrievable annotations were notably two odorant binding proteins (for R-C/C-S comparisons, respectively: AARA007908: FCs=5.17 and 0.17; AARA004722: FCs=3.24 and 0.19).

Significant differential expression of some members of the detoxification gene families associated with metabolic resistance were observed among R-S and C-S comparisons (Table 2; Figure 3A). These included nine CYP450s (CYP9K1, CYP9J5, CYP6AA1, CYP4C36, CYP6AA1, CYP9L1, CYP6M2, CYP6M3 and CYP6P4), six GSTs (GSTE2, GSTE3, GSTE4, GSTE5, GSTE7 and GSTD3) and two COEs (AARA016305 and AARA016468). With the exception of GSTD3 and GSTE3, the FCs of all of these detoxification enzymes increased in response to malathion exposure (Table 2). Two additional CYP450s were also upregulated between R-C conditions (CYP4G16, FC=3.40; and CYP4G17, FC=2.03) (Supplementary Figure S1; Table 2).

**Table 2.**
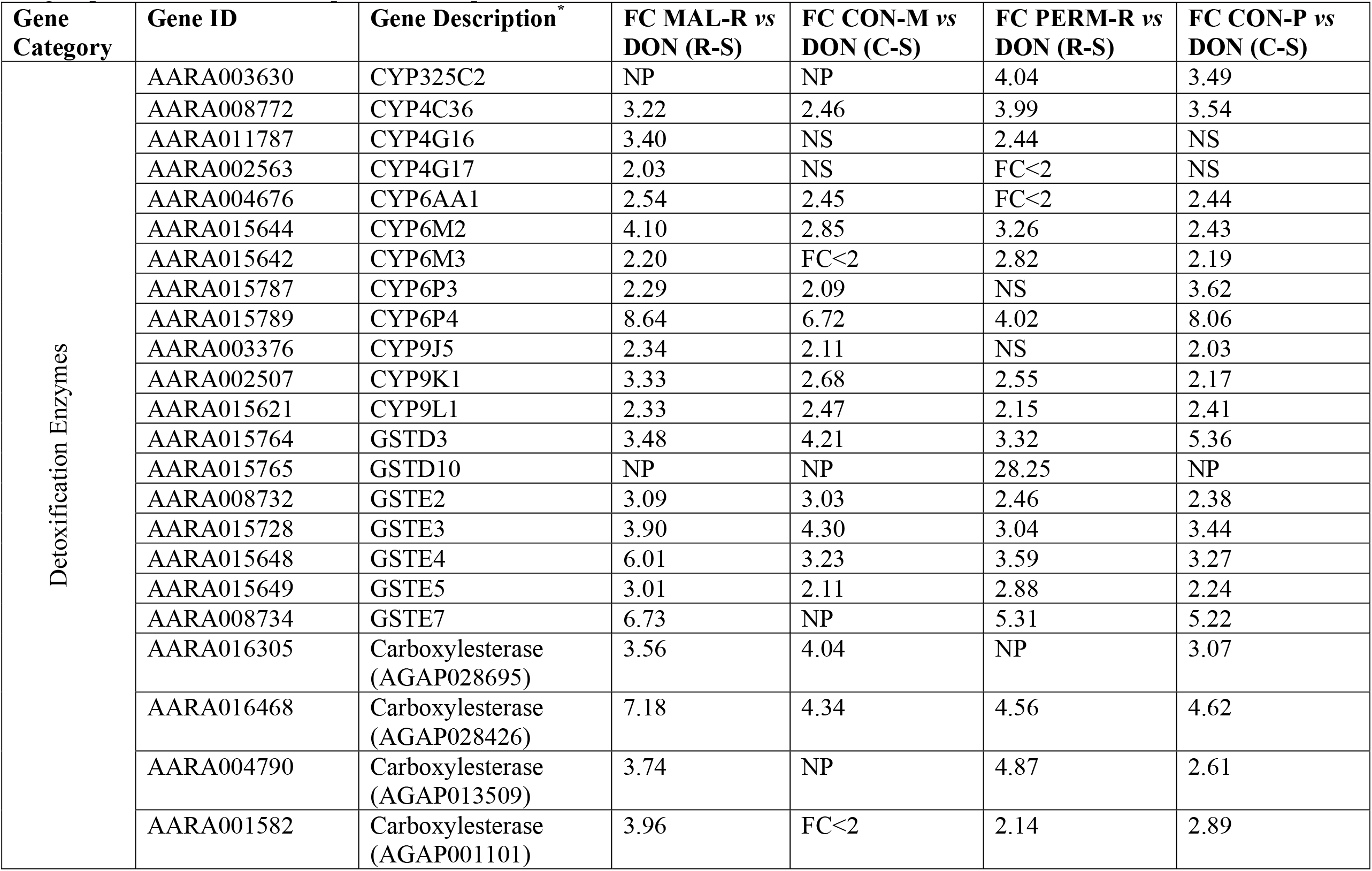
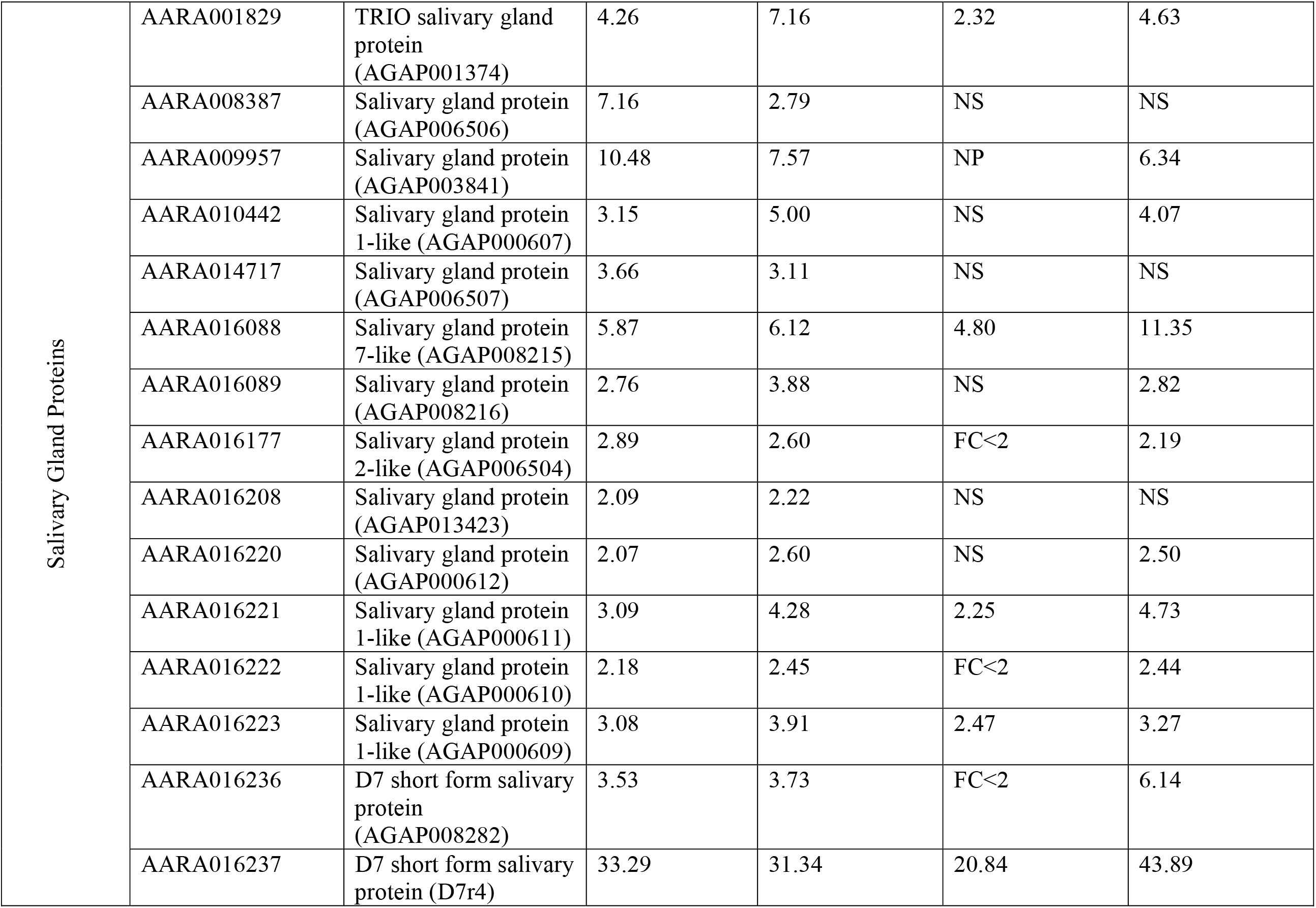
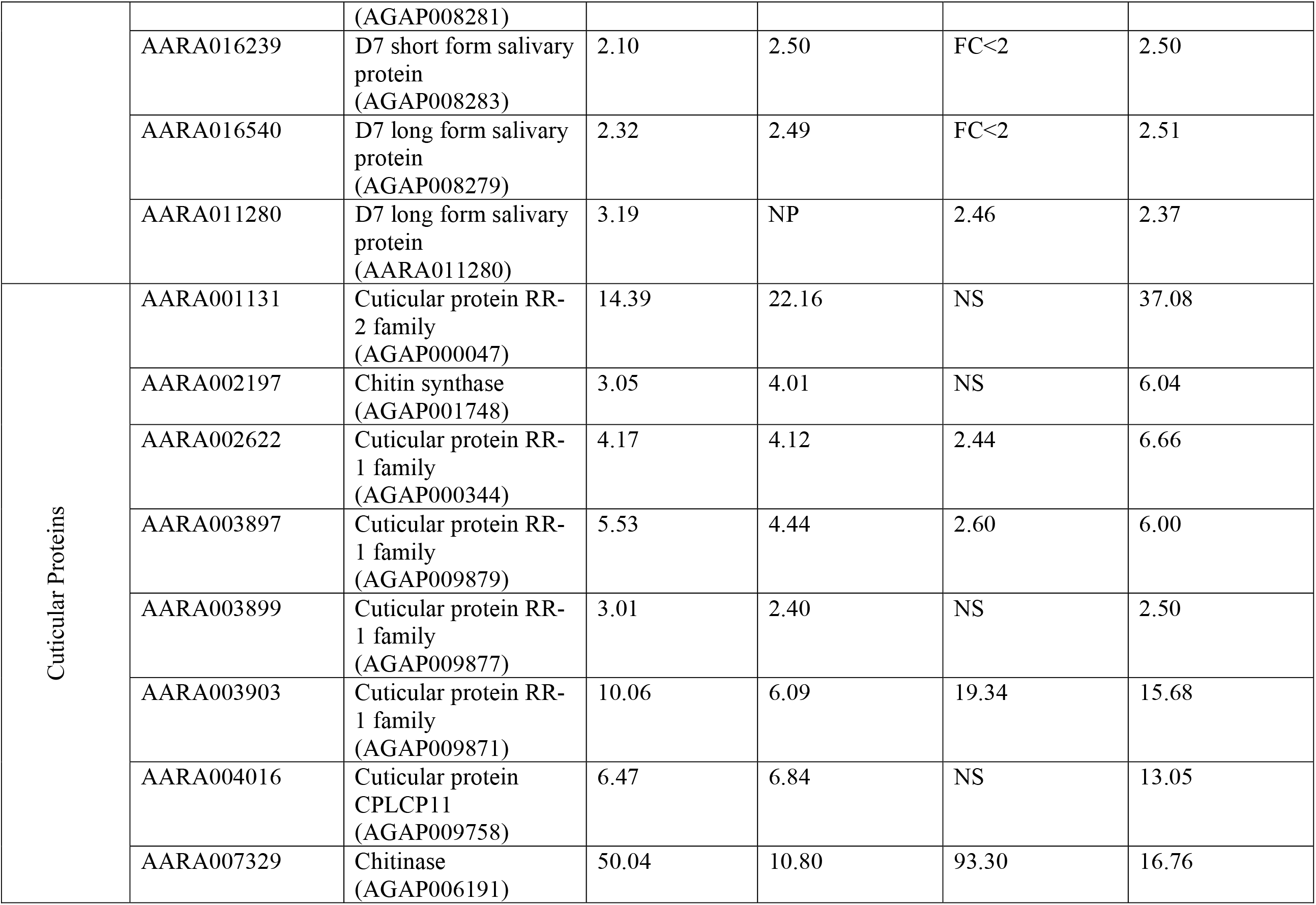
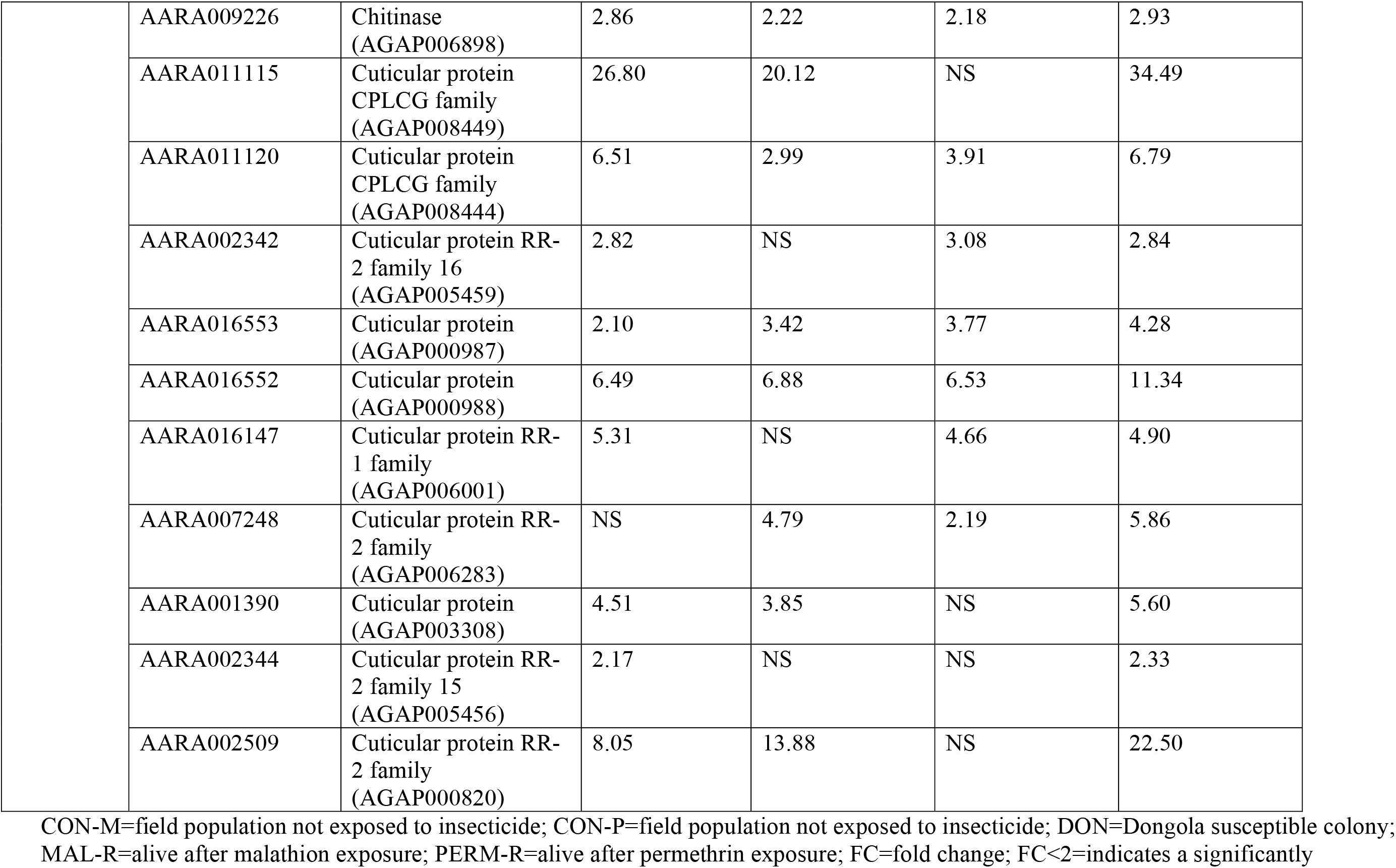
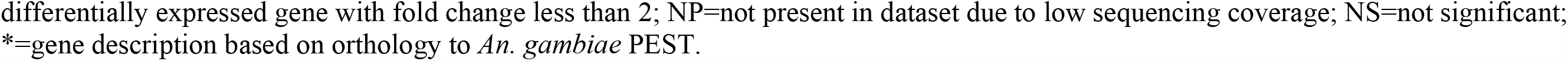
Significantly differentially expressed genes of interest in comparisons of resistant *vs* susceptible (R-S) and control *vs* susceptible (C-S) groups in the malathion and permethrin experiments (FDR<0.05 and FC>2).

**Figure 3.**
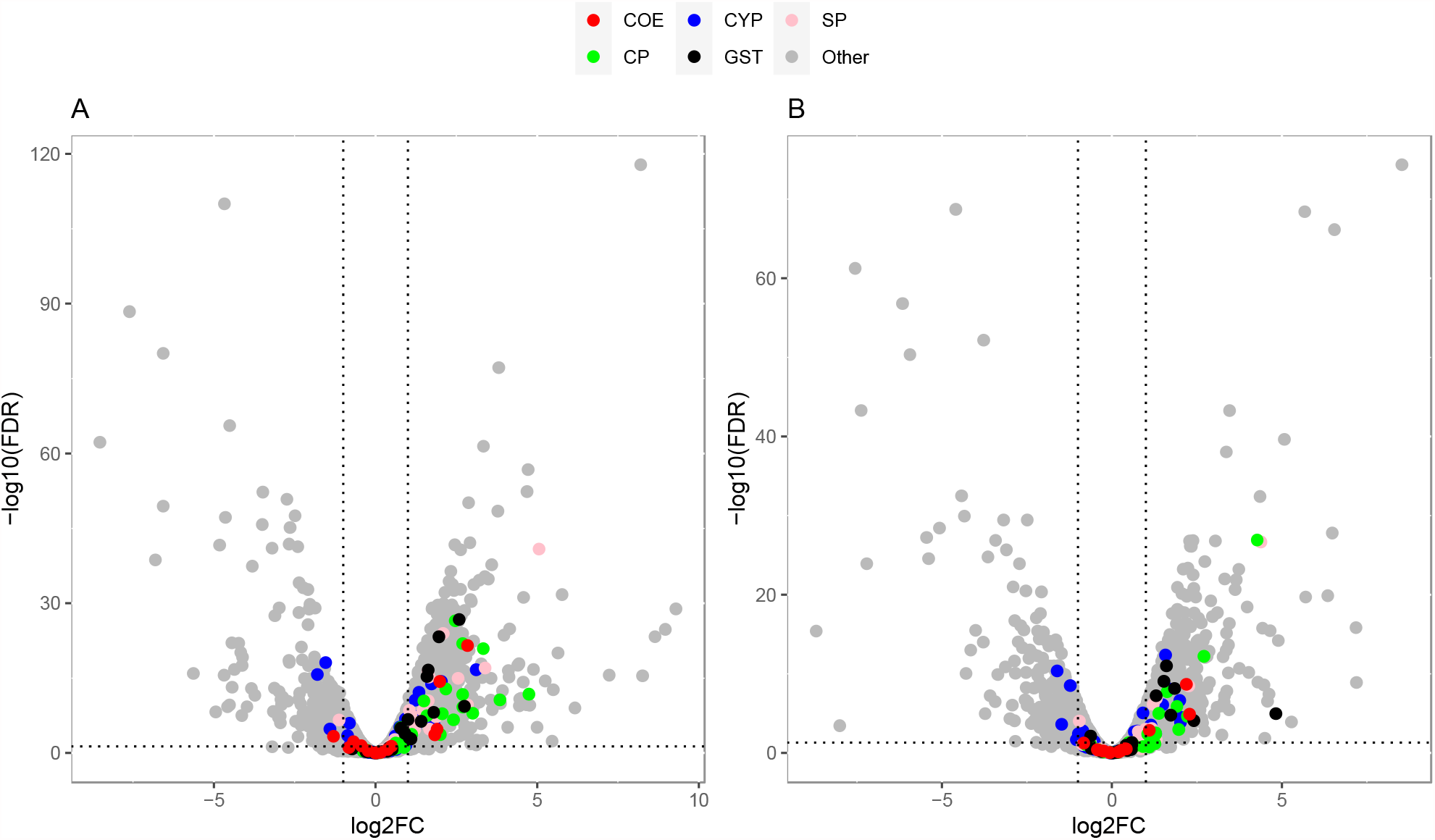
Volcano plots of gene expression for MAL-R vs DON (A) and PERM-R vs DON (B). Detoxification gene families are indicated in red (COE: carboxylesterases), blue (CYP: cytochrome P450s) and black (GST: glutathione-S-transferases). Cuticular proteins are indicated in green (CP) and salivary gland proteins are indicated in pink (SP). In each plot, genes overexpressed in the population are >0 on the x-axis. Vertical dotted lines indicate 2-fold expression differences and the horizontal line indicates a *P*-value of 0.01.

Significant differential expression of eighteen mosquito salivary gland proteins were identified among R-S and C-S comparisons (Table 2; Figure 3A), most notably D7r4 short form salivary protein (FCs=33.29 and 31.34 for R-S/C-S, respectively), TRIO salivary gland protein (FCs=4.26 and 7.16), AARA009957 (FCs=10.48 and 7.57) and salivary gland protein 7 (FCs=5.87 and 6.12). Among these salivary gland proteins, twelve were downregulated following malathion exposure (Table 2); one salivary gland protein was significantly overexpressed between R-C conditions (AARA008387, FC=2.04). Furthermore, fifteen proteins associated with cuticular function were significantly overexpressed in the R-S condition, including chitinase (AARA007329) (FCs=50.04 and 10.80 for R-S/C-S, respectively), cuticular protein CPLCG family (AARA011115) (FCs=26.80 and 20.12), cuticular protein RR-2 family (AARA001131) (FCs=14.39 and 22.16) and cuticular protein RR-1 family (AARA003903) (FCs=10.06 and 6.09). The majority of these were upregulated after insecticide treatment (Table 2), with an additional cuticular protein RR-2 family member, significantly overexpressed between R-C conditions (AARA017766, FC=2.45) (Supplementary Figure S1).

In malathion resistant mosquitoes, several ontologies were enriched in genes overexpressed relative to susceptible mosquitoes (Table S3). In particular, many of these ontologies were associated with metabolic processes, including “cellular metabolic process” (GO:0044237), “catalytic activity” (GO:0003824) and “generation of precursor metabolites and energy” (GO:0006091). Between R-C conditions, additional metabolic ontologies were upregulated, including “generation of precursor metabolites and energy” (GO:0006091) and “cellular metabolic process” (GO:0044237), potentially associated with increased physiological stress in response to insecticide exposure (21).

### Differentially expressed genes associated with permethrin resistance

Differential transcription analysis for the permethrin experiment was performed relative to both DON and Sekoru (SEK) susceptible laboratory strains; the latter analysis was performed with the assumption that this more geographically proximate colony from Ethiopia would be a better biologically comparator than DON. However, greater variation in gene expression was observed, with 2183 (23.5%; 1057 upregulated and 1126 downregulated) and 3412 (35.0%; 1153 upregulated and 2259 downregulated) genes significantly differentially expressed between SEK and mosquitoes that survived permethrin exposure and non-exposed field mosquitoes, respectively (Supplementary Figure S2; Table 1). A multi-dimensional scaling plot revealed significant variation between SEK and all other mosquito populations (Supplementary Figure S3); downstream analyses for the permethrin experiment were therefore performed relative to DON.

Consistent with the malathion experiment, three pairwise comparisons were conducted for permethrin: resistant *vs* susceptible (R-S; PERM-R *vs* DON), resistant *vs* unexposed control (R-C; PERM-R *vs* CON-P) and unexposed control *vs* susceptible (C-S; CON-P *vs* DON). Among mosquitoes that survived permethrin exposure and non-exposed field mosquitoes, 1074 (10.9%; 673 upregulated and 401 downregulated) and 889 (8.9%; 594 upregulated and 295 downregulated) genes were significantly differentially expressed (at *P*-values adjusted for multiple testing based on a FDR<0.01 and FC>2), respectively, when compared to the susceptible Dongola strain (Figure 2B; Table 1). A total of 334 (3.5%; 179 upregulated and 155 downregulated) genes were significantly differentially expressed in permethrin survivors as compared to their non-exposed counterparts (Figure 2B; Table 1).

Of the genes that were differentially expressed in all treatment groups (n=35), 9 were upregulated while 14 were downregulated in one or more conditions (Figure 2B). Eleven had retrievable annotations, which were mostly molecular functions or biological processes (for R-C/R-S/C-S comparisons: AARA015710=CLIP-domain serine protease, FCs=2.21, 4.35 and 1.97; AARA015772=cytoplasmic actin, FCs=4.24, 51.86 and 12.20; AARA016057=ATP binding cassette transporter, FCs=0.41, 2.39 and 5.82; AARA016221=salivary gland protein 1-like, FCs=9.47, 2.25 and 4.73; AARA002374=MIP18 family protein CG7949, FCs=2.38, 4.01 and 1.67; AARA003468=peptide methionine sulfoxide reductase, FCs=3.63, 0.42 and 0.11; AARA003599=TRPL translocation defect protein 14 isoform, FCs=2.20, 3.47 and 1.57; AARA009096=diacylglycerol kinase 1 isoform, FCs=0.41, 0.22 and 0.53; AARA016129=sorbitol dehydrogenase, FCs=0.04, 0.35 and 7.99; AARA017544=serine protease 7-like, FCs=2.58, 4.70 and 1.82; and AARA018460=lysosomal alpha-mannosidase, FCs=0.42, 4.24 and 10.04, respectively).

A total of 436 genes were differentially expressed commonly in the R-S and C-S groups (Figure 2B). The top 10 over-expressed genes with retrievable annotations were similar to the malathion experiment, including chitinase (AARA007329: FCs=93.30 and 16.76 for R-S/C-S comparisons, respectively), D7r4 short form salivary protein (AARA016237: FCs=20.84 and 43.89), cytoplasmic actin (AARA015772: FCs=51.86 and 12.20), alkaline phosphatase (AARA002132: FCs=29.70 and 13.74), sulfotransferase (AARA016556: FCs=33.61 and 16.61), polyubiquitin (AARA016579: FCs=21.57 and 67.65) and ADP/ATP carrier protein (AARA017958: FCs=25.15 and 10.50). Cuticular protein RR-1 (AARA003903: FCs=19.34 and 15.68) and hexamerin (AARA016988: FCs=15.78 and 7.50) were also highly upregulated.

Consistent with the malathion experiment, key metabolic enzymes were significantly differentially expressed between R-S and C-S comparisons (Table 2; Figure 3B), including eight CYP450s (CYP6M2, CYP4C36, CYP6AA1, CYP9K1, CYP6M3, CYP6P4, CYP325C2 and CYP9L1), six GSTs (GSTE2, GSTE3, GSTE4, GSTE5, GSTE7 and GSTD3) and three COEs (AARA016468, AARA001582 and AARA004790). Six of these detoxification genes were downregulated following permethrin exposure, including CYP6AA1, CYP9L1, GSTD3, GSTE3 and two COEs (AARA016468 and AARA001582). One additional CYP450 was also significantly overexpressed between R-C conditions (CYP6Z3, FC=2.02). A further GST (GSTD10) was highly overexpressed in both R-S and R-C conditions (FCs=28.25 and 5.94, respectively), but was not present at sufficient sequence coverage in the C-S comparison. In addition, six mosquito salivary gland proteins were identified among R-S and C-S comparisons (Table 2), most notably D7r4 short form salivary protein (FCs=20.84 and 43.89 for R-S/C-S, respectively), and salivary gland protein 7 (FCs=4.80 and 11.35), which in contrast to the malathion experiment, were both downregulated in response to permethrin exposure. A further eleven proteins associated with cuticular function displayed differential expression patterns (Table 2; Figure 3B), including chitinase (AARA007329; FCs=93.30 and 16.76, for R-S/C-S, respectively), cuticular protein RR-1 family (FCs=19.34 and 15.68,) and cuticular protein (FCs=6.53 and 11.34, for R-S/C-S). An additional chitinase was significantly overexpressed between R-C conditions (AARA007329, FC=5.56) (Supplementary Figure S1).

Similar to the malathion experiment, ontologies enriched in the permethrin experiment also included terms related to “metabolic process” (GO:0008152), “generation of precursor metabolites and energy” (GO:0006091), “oxidoreductase activity” (GO:0016491) and “carbohydrate metabolic process” (GO:0005975).

### Differentially expressed genes associated with multi-insecticide resistance

A total of 717 (45.7%; 512 upregulated and 205 downregulated) transcripts were significantly differentially expressed in mosquitoes that survived either malathion or permethrin exposure, compared to the susceptible strain (Table S5). Eight key upregulated metabolic enzymes were shared between both resistant groups (MAL-R *vs* DON and PERM-R *vs* DON), including six CYP450s (CYP6P4, CYP4C36, CYP4G16, CYP6M3 and CYP9K1 and CYP9L1), six GSTs (GSTD3, GSTE2, GSTE3, GSTE4, GSTE5 and GSTE7) and three COEs (AARA004790, AARA016468 and AARA001582) (Table 2; Figure 4); two additional CYP450s (CYP9M2 and CYP304B1) were both downregulated. Unique detoxification DEGs to the malathion resistant group were CYP9J5 (FCs=2.34 and 2.11 for R-S/C-S, respectively), CYP6P3 (FCs=2.29 and 2.09) and one COE (AARA016305: FCs=3.56 and 4.04 for R-S/C-S, respectively). One detoxification DEG was unique to the permethrin resistant population, CYP325C2 (FCs=4.04 and 3.49, for R-S/C-S, respectively), but was not present at sufficient sequence coverage in the malathion resistant population.

**Figure 4.**
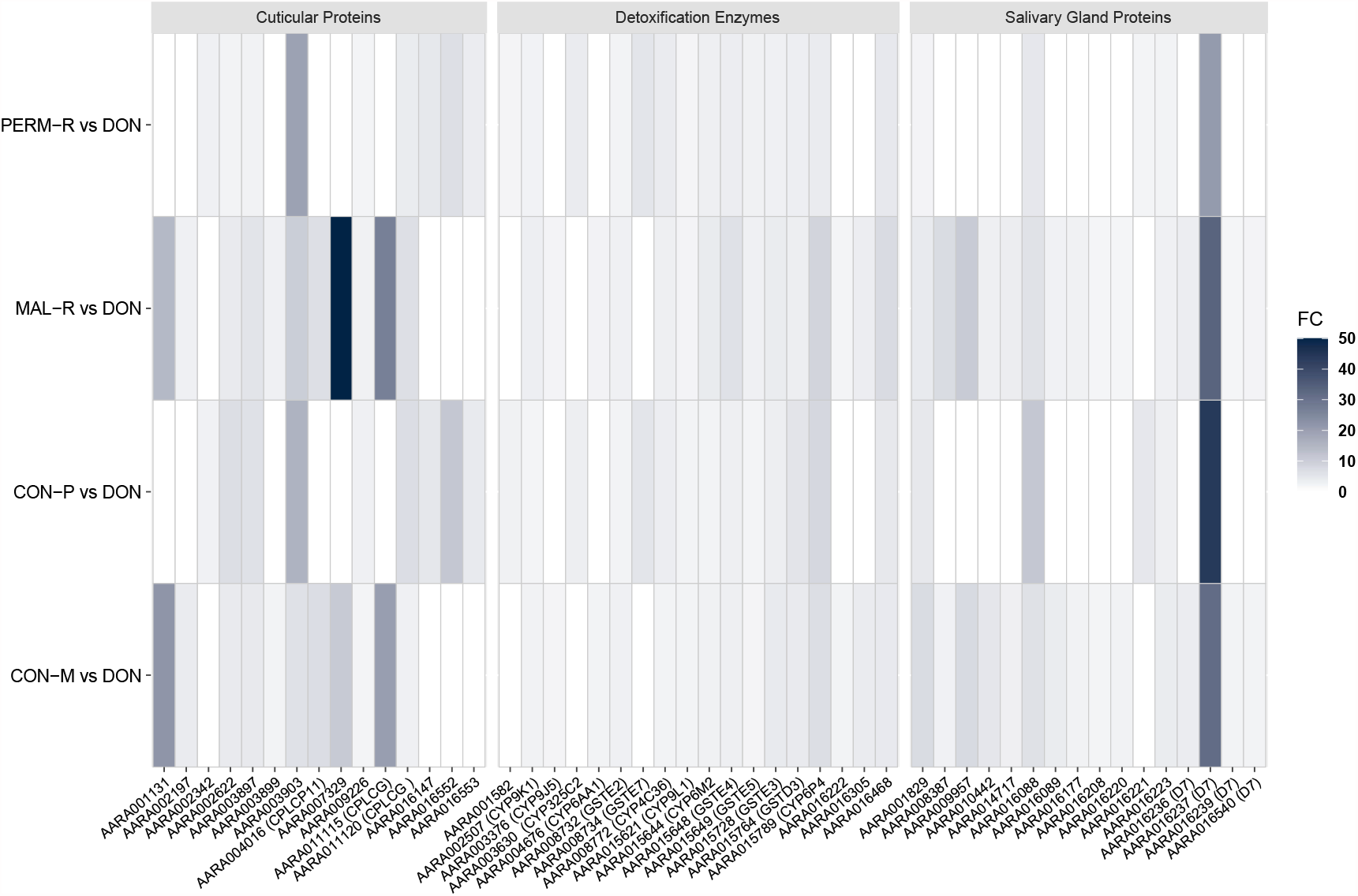
Heatmaps summarizing expression of cuticular proteins, detoxification enzymes and salivary gland proteins, showing fold-change values relative to the susceptible strain. CON-M=field population not exposed to malathion; CON-P=field population not exposed to permethrin; DON=Dongola susceptible colony; MAL-R=alive after malathion exposure; PERM-R=alive after permethrin exposure; FC=fold change.

Among salivary gland DEGs, six were shared between both resistant populations (Table 2; Figure 4): D7r4 short form salivary protein (AARA016237), D7 long form salivary gland protein (AARA011280), salivary gland protein 1-like members (AARA016223 and AARA016221), TRIO salivary gland protein (AARA001829) and salivary gland protein 7-like members (AARA016088). Twelve additional salivary gland proteins were exclusive to the malathion resistant population and none to the permethrin resistant population (Table 2; Figure 4).

Among cuticular DEGs, ten were shared between both resistant populations: cuticular protein RR-1 family members (AARA002622, AARA003897, AARA003903 and AARA016147), chitinases (AARA007329 and AARA009226), a cuticular protein CPLCG family member (AARA011120), a cuticular protein RR-2 family 16 member (AARA002342), cuticular proteins (AARA016553 and AARA016552). There were eight and one DEGs which were unique to the malathion and permethrin resistant populations, respectively (Table 2; Figure 4).

Finally, we mined the RNA-seq data to investigate expression patterns of other recently described resistance mechanisms in *An. gambiae* complex members (12,15) (Table 3). We identified orthologues in *An. arabiensis* of four α-crystallins, two hexamerins, ATPase subunit e and SAP2 which were significantly differentially expressed between R-S/C-S conditions.

**Table 3.**
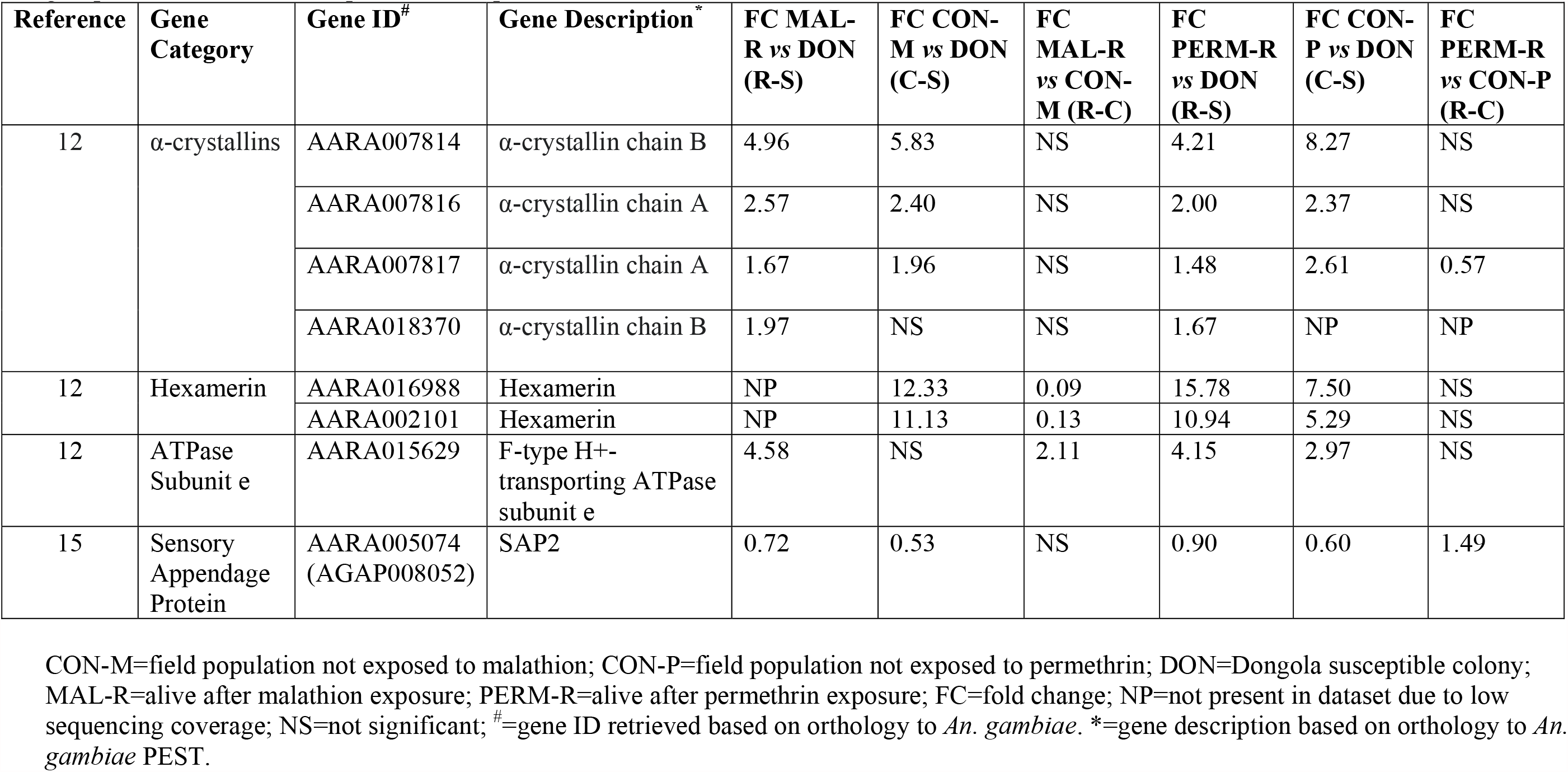
Significantly differentially expressed genes of interest in comparisons of resistant *vs* susceptible (R-S) and control *vs* susceptible (C-S) groups in the malathion and permethrin experiments (FDR <0.05).

### Detection of resistance target site mutations

RNA-Seq reads from the malathion and permethrin experiments were screened for target site mutations associated with DDT, pyrethroid, organophosphate or carbamate resistance and known voltage-gated sodium channel (VGSC) mutations in *An. gambiae* s.l. (Tables S6 and S7). Consistent with the target site PCR data generated in this study, we did not detect the presence of either L1014S *kdr* or G119S *Ace-1* mutations in any populations. The L1014F-*kdr* mutation was detected in all groups except DON, with average population allele frequencies of CON-M=27%; CON-P=24%; MAL-R=31%; PERM-R=79%; and SEK=55% (Table S6). None of the previously described GSTe2 target site mutations (L119F and I114T) (23,24) were present in our dataset, nor was N1575Y, which is linked to L1014F-*kdr* and found at variable frequencies in parts of West and Central Africa (25-27). Of 20 recently described non-synonymous VGSC mutations from West and Central Africa (28), we detected the presence of seven (R254K, A1125V, I1868T, P1874L, F1920S, A1934V and I1940T) across the Asendabo field population at very low frequencies (range of 1-7%); 2 of these were also found in SEK (I1868T and I1940T).

### qRT-PCR validation of relative expression levels estimated by RNA-Seq

Quantitative RT-PCR was used to validate the FCs of eleven genes (CYP4G16, CYP4G17, GSTM3, CPR130, GSTE7, CYP6M2, D7r4 short form salivary protein, chitinase, cuticular protein RR-1 family, CYP6M3 and GSTE3), relative to two housekeeping genes (40S ribosomal protein S7; RPS7 and ubiquitin) (Figure 5). The majority of the qRT-PCR results supported the directionality of the changes in expression levels as estimated by RNA-Seq.

**Figure 5.**
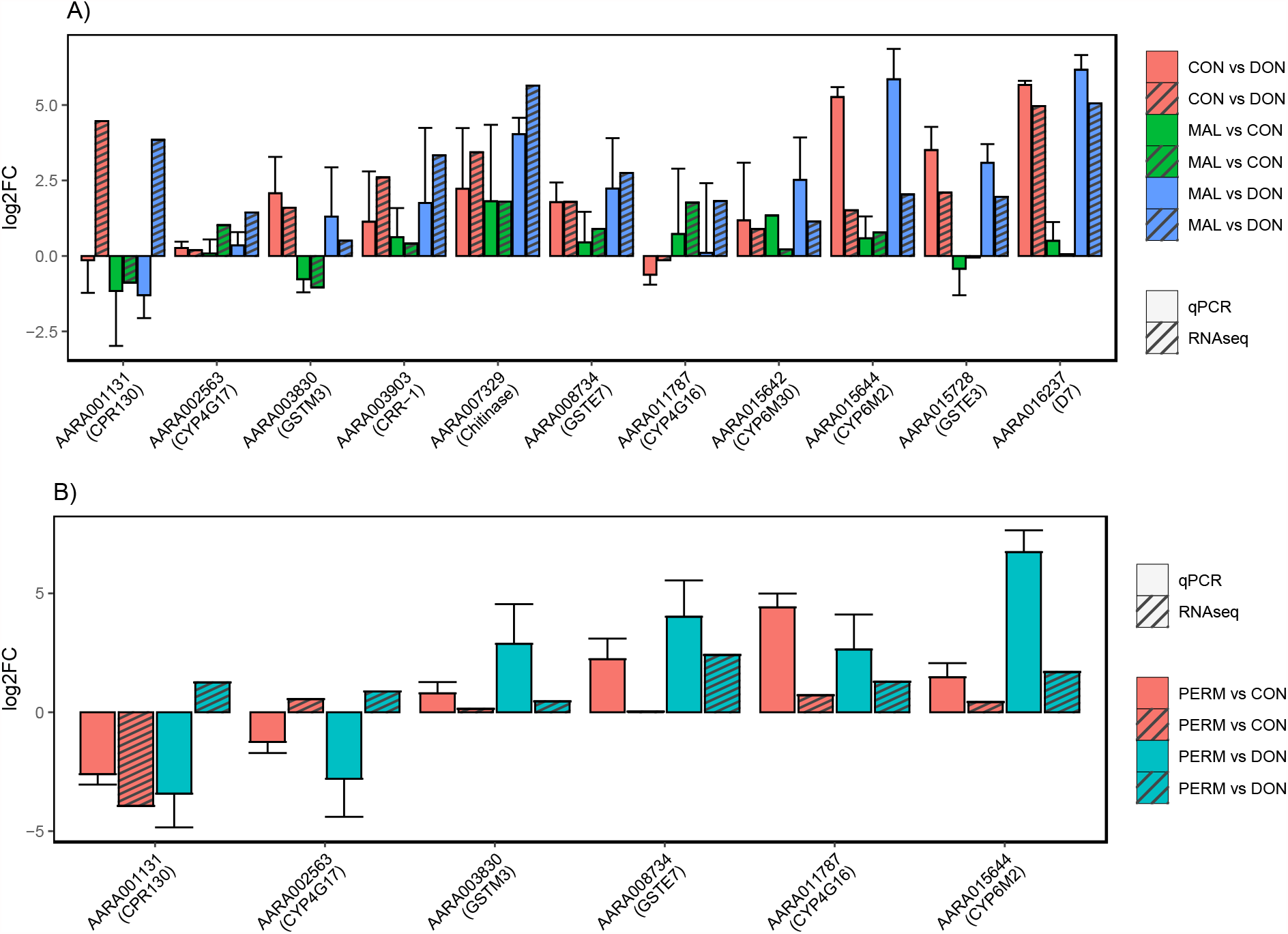
Comparison of expression levels of DEGs measured by qRT-PCR and RNA-Seq in malathion (A) and permethrin (B) experiments.

## Discussion

Using a whole transcriptomic approach, we investigated mechanisms conferring resistance to malathion and permethrin in *An. arabiensis* from southwest Ethiopia. Our analyses allowed for comparisons between insecticides, to detect shared expression patterns between different active ingredients and to identify novel diagnostic markers associated with phenotypic resistance. In addition to malathion and permethrin resistance, the field population was also resistant to deltamethrin but remained susceptible to alpha-cypermethrin, bendiocarb and propoxur. A previous study from the same region reported susceptibility to the putative diagnostic doses of clothianidin (neonicotinoid) and chlorfenapyr (pyrrole) (29). Bioassay results indicated that insecticide-specific mechanisms may be important in this *An. arabiensis* population, as demonstrated by the lack of cross-resistance between active ingredients belonging to the same chemical class (for example permethrin and alpha-cypermethrin). Insecticide resistance profiles in Asendabo aligned with recent nationwide insecticide resistance monitoring results (3). However, bendiocarb and alpha-cypermethrin tolerance appeared dynamic, with reduced local *An. arabiensis* mortality to both chemicals reported during previous years but absent in our study (3).

In both malathion and permethrin resistant groups, several CYP450s and GSTs, which have been associated with pyrethroid and DDT resistance in populations of *An. arabiensis*, were overexpressed. Upregulation of CYP6M2, CYP6M3, CYP6P4, CYP9K1 and GSTE4, which were shared between both resistant groups, has been documented in pyrethroid and DDT resistant *An. arabiensis* from central Sudan (30). In addition, we detected overexpression of CYP4C36, CYP6AA1, CYP9L1, GSTD3, GSTE2, GSTE3, GSTE5, GSTE7 and three carboxylesterases (AARA016468, AARA004790 and AARA001582) in both resistant groups; carboxylesterases have previously been implicated in pyrethroid resistance in *An. funestus* from Malawi (30). Overexpression of CYP6P3 and CYP9J5, which were exclusive to malathion survivors, has also been observed in permethrin-resistant *An. arabiensis* from Sudan (31). Many of these CYP450s are regularly reported from multi-insecticide resistant populations of *An. gambiae* across sub-Saharan Africa and have been shown to metabolize different combinations of type I and type II pyrethroids, DDT and pyriproxyfen *in vitro* (7-9, 32-34; reviewed by 35). *In vivo* functional characterization of CYP6M2 and CYP6P3 in *An. gambiae* demonstrated that overexpression enhanced susceptibility to malathion by catalysing the bioactivation of this insecticide to its more toxic metabolite malaoxon by a CYP450-mediated mechanism (36); with CYP6M2 increasing malaoxon production to a greater degree compared to CYP6P3 (37). Our contradicting results may be explained by the relative activity of the transcription factor Maf-S, which when knocked-down has been shown to increase survival to malathion exposure (13) and was not found to be significantly differentially expressed in this study. CYP325C2, which was the only unique CYP450 overexpressed in our permethrin resistant population, has been reported from *An. arabiensis* in Kenya (38) and Cameroon (39) with reduced susceptibility to deltamethrin. Interestingly, CYP325C2 was not identified as a DEG in previous transcriptomic analysis performed among deltamethrin and DDT survivors from Asendabo, which may indicate that it is specific to permethrin resistance in this field population (17). Following permethrin exposure, CYP6Z3 was also significantly upregulated in survivors compared to the unexposed population; overexpression of this enzyme has also been implicated in lambda-cyhalothrin resistance in *An. arabiensis* from Tanzania (40).

In Ethiopia, spatial and temporal patterns of insecticide resistance have generally correlated with changes in national malaria vector control policy. Intense pyrethroid resistance is not unexpected given the quantity of conventional LLINs which have been distributed across the region (>80 million since 2008), while the continued presence of malathion resistance is more surprising. Malathion was last used extensively for malaria control from 2003 to 2005 by the NMCP in areas with reported DDT resistance (41,42). Between 2005-2017, malathion susceptibility was monitored in 127 sentinel sites in Ethiopia, with evidence for possible resistance at 55 sites, confirmed resistance at 36 sites and susceptibility at 36 sites (reviewed by 43). In general, resistance instability in the absence of insecticidal pressure, largely attributable to fitness costs, has been well documented among a number of medically-important vector species (44-46); with some notable exceptions, particularly dieldrin resistance (47). Selection experiments using field populations of *An. gambiae* have determined that the rate of resistance decay to full pyrethroid susceptibility from moderate resistance intensity can be as little as 15 generations or approximately 1.3 years in typical African settings (48). Our transcriptome data revealed shared overexpression of detoxification enzymes between malathion and permethrin resistant groups, which may be responsible for cross-resistance due to ongoing pyrethroid selection and as a result, maintaining decreased malathion susceptibility.

Another explanation for the continued persistence of malathion resistance in this field population might be that underlying resistance mechanisms impart other physiological benefits to individuals in both the presence and absence of insecticidal exposure. We identified nineteen cuticular proteins and associated enzymes which in some cases were upregulated by more than fifty- or ninety-fold in resistant groups compared to the susceptible strain. These were generally much more highly overexpressed than any of the detoxification enzymes and some of which were observably induced by insecticide exposure (including cuticular protein RR-1 family; AARA003903, chitinase; AARA007329 and cuticular protein CPLCG; AARA011115). Evidence is emerging to strongly support a key role for cuticular thickening as a generalist mechanism of insecticide resistance across *Anopheles* populations, through either enriched deposition of cuticular hydrocarbons or changes to structural components of the procuticle (49). Thicker femur cuticles can delay the penetration rate of contact chemicals and/or increase the time available for metabolic processes to inactivate the insecticide before it causes inhibition, thus potentially producing a more intense, non-specific resistance phenotype (50). Following malathion exposure, our field population was characterized by a significant increase in CYP4G16 and CYP4G17 expression; both genes are known to facilitate hydrocarbon production, with the former catalysing epicuticular hydrocarbon biosynthesis (50,51). Previous analyses of the Asendabo population also support the potential involvement of cuticular resistance, via increased cuticular hydrocarbon quantities but not procuticle thickness (17). Recent multiplex qRT-PCR assays have been developed with CYP4G16 as a candidate surveillance marker for metabolic resistance in *An. gambiae* which will begin to improve our understanding of its relative involvement in regional cuticular resistance (52). Among the genes we selected for qPCR validation, chitinase (AARA007329) was very highly overexpressed, induced by exposure to malathion (FCs=50.04 and 10.80, for R-S/C-S, respectively) and permethrin (FCs=93.3 and 16.76) and may represent an informative cuticular-associated gene for resistance monitoring in *An. arabiensis* populations. Further investigation is required to determine whether chitinase overexpression is a causative factor in resistance or if it is closely associated with a resistance-conferring variant, as it might be expected to enhance insecticide toxicity by promoting faster cuticle degradation (53).

In this study, we also detected another putative resistance mechanism in the form of eighteen upregulated salivary gland proteins, particularly the D7 short form salivary protein (the orthologue of D7r4 in *An. gambiae*), which was overexpressed by more than twenty-to thirty-fold following malathion exposure but was notably downregulated following permethrin exposure. Overexpression of D7r4 has been observed in pyrethroid-resistant *An. arabiensis* populations from Sudan, Uganda and Zanzibar (31,40,54) and carbamate- and pyrethroid-resistant *An. funestus* and *An. gambiae* (14,55); this is the first report of D7r4 associated with organophosphate (malathion) resistance. It has been suggested that D7 overexpression is symptomatic of a disruption in the tissue-specificity of these salivary gland proteins, allowing these proteins to interact with insecticides in tissues other than the salivary glands (14). Furthermore, *in silico* modelling of the protein structure of D7r4 has shown it can accommodate bendiocarb in its central binding pocket, supporting a role for this molecule in binding and sequestering insecticide or insecticide metabolites, rather than by direct detoxification (14). Similarly, we detected overexpressed candidate α-crystallins, hexamerins and an ATPase subunit which have been proposed to play as yet undefined functions in binding and sequestering insecticides (12). By comparison to *An. gambiae*, our understanding of resistance mechanisms in *An. arabiensis* is far more limited; however, our findings highlight several potential shared pathways between these major vector species that warrant further investigation.

In addition to gene expression patterns, we also investigated the prevalence of known resistance target site mutations in our field population. We detected L1014F-*kdr* at moderate to high allele frequencies among permethrin survivors, but with little evidence for ongoing selection at this locus. We also confirmed the absence of L1014S-*kdr*, N1575Y, G119S-*Ace-1* and two GSTe2 mutations (L119F and I114T) (23-25), which have yet to be reported in Ethiopia (3,4,17). Furthermore, from our RNA-Seq data, we detected the presence of seven novel mutations in the VGSC of our pooled *An. arabiensis* populations; one in domain one (in the linker between transmembrane segments four and five; R254K), one in the linker between domains two and three (A1125V) and five in the internal carboxyl tail (I1868T, P1874L, F1920S, A1934V and I1940T). These belong to a group of 14 non-synonymous substitutions in the VGSC recently described in *An. gambiae* and *An. coluzzii*, which have likely evolved in association with L1014F-*kdr* and appear to have been positively selected following decades of DDT/pyrethroid use (28). In particular, the substitutions located in the C-terminal tail have been proposed to disrupt the confirmation of the DIII-DIV linker subdomain, which is normally bound in close proximity to the DIV S6 helix, inactivating the VGSC (28). The expected outcome would be altered channel inactivation, but this awaits functional validation.

## Conclusions

Using a whole transcriptomic approach, we investigated resistance mechanisms to two public health insecticides in *An. arabiensis* from Ethiopia. Study findings revealed shared detoxification enzymes between organophosphate and pyrethroid-resistant vectors and highly overexpressed cuticular-associated proteins and salivary gland proteins, which may play a role in enhancing vector resistance. The advantages of adopting a transcriptomic approach are evidenced by its universal mechanistic characterization, allowing for the discovery of novel candidate resistance genes, which warrant functional validation to determine their contributions to insecticide resistance, including their potential to confer cross-resistance between different insecticides with the same mode of action.

## Materials and Methods

### Study area and mosquito collections

Adult mosquitoes were collected from Asendabo, Oromia region, Ethiopia (7°40’31” N, 36°52’56” E), where organophosphate and pyrethroid resistance had been previously reported in *An. arabiensis* populations (3). Mosquitoes were sampled at the end of the long rainy season, between 3^rd^ September-10^th^ October 2017, following IRS with bendiocarb by the National Malaria Control Program (NMCP) in this area in June 2017.

Upon obtaining householder consent, indoor-resting, blood-fed female *Anopheles* mosquitoes were collected from the walls of 12 houses (situated approximately <5km apart) between 4:00 and 6:00 am using handheld aspirators. Mosquitoes were held in paper cups with access to 10% sucrose and transported to the Tropical and Infectious Diseases Research Center (TIDRC) in Sekoru, Oromia region (7°54’50” N, 37°25’23.6” E). F_1_ progeny were obtained from field-collected mosquitoes using forced-oviposition (18). Blood-fed, field-collected mosquitoes, morphologically identified as *An. gambiae* s.l. (56), were maintained for 4-5 days until fully gravid and checked daily for survival. Each fully gravid female was transferred to a 1.5 ml microcentrifuge tube containing damp cotton wool and allowed to lay eggs. Eggs from 246 adult *An. gambiae* s.l. were transported to the U.S. Centers for Disease Control and Prevention (CDC), Atlanta, USA, and pooled for rearing in the CDC insectary.

*An. arabiensis* from the insecticide susceptible Dongola reference strain (originating from Sudan, obtained from the Malaria Research and Reference Reagent Resource Center, MR4) and the Sekoru insecticide susceptible laboratory strain (originating from Ethiopia, obtained from the Vector Biology and Control Research Unit, TIDRC, Jimma University) (57), were also reared in the CDC insectaries. All adult mosquitoes were maintained under standard insectary conditions (27±2°C, 80% relative humidity, light:dark cycles of 14:10 hours) with access to 10% sucrose solution *ad libitum*. F_1_ adult females of each strain were randomly mixed in cages for subsequent bioassays.

### Insecticide resistance bioassays

CDC bottle bioassays for malathion (organophosphate) and permethrin (pyrethroid) were conducted according to published guidelines (20). Stock solutions of the diagnostic dose required to kill 100% of susceptible mosquitoes (malathion: 50μg/bottle and permethrin: 21.5μg/bottle), were prepared by diluting technical grade insecticide in 50ml of acetone. Each Wheaton 250ml glass bottle along with its cap was coated with 1ml of the stock solution by rolling and inverting the bottles. In each test, a control bottle was coated with 1 ml of acetone. Bottles were left to dry in the dark for three hours and were washed thoroughly and re-coated before every test. Following a two-hour acclimatization period in paper cups with access to 10% sucrose, approximately, 20-25 unfed, 3 day-old adult female *An. gambiae* s.l. were introduced into each bottle using a mouth aspirator and knock-down/mortality was recorded after 30 minutes of exposure. Additionally, a susceptible reference *An. arabiensis* strain (Dongola or Sekoru) was assayed in parallel. Bioassays were conducted between 15:00 and 17:00 each day to avoid any bias in RNA transcript expression related to circadian rhythm. Multiple replicates were performed per insecticide to obtain sufficient phenotyped material for RNA-sequencing analysis. A mosquito was defined as ‘alive’ at the diagnostic time if it was capable of standing and flying in a coordinated manner; surviving mosquitoes (defined as resistant) and non-exposed mosquitoes (from acetone-treated bottles) were stored separately at -80°C. Additionally, non-exposed, unfed, 3 day-old adult female *An. arabiensis* from the Sekoru and Dongola susceptible laboratory strains were also preserved for analysis at -80°C.

Additional resistance intensity bioassays were undertaken with F_1_ field mosquitoes to characterize susceptibility levels to carbamates (bendiocarb and propoxur) and pyrethroids (alpha-cypermethrin, deltamethrin and permethrin), following exposure to 1, 2, 5 and 10 times the diagnostic doses. Bioassay data were interpreted according to the WHO criteria: mortality of 98% or higher indicates susceptibility, mortality of 90-97% is suggestive of resistance, and mortality of less than 90% indicates resistance (58). Mortality in untreated control bottles was less than 5% in all resistance intensity bioassays. Mean percent mosquito mortality was calculated across all replicates for a given insecticide.

### Molecular species identification

Prior to pooling specimens for RNA extraction, 4-6 legs from each mosquito tested in bioassays were removed and genomic DNA was extracted using the Extracta™ DNA Prep for PCR-Tissue kit (QuantaBio, USA), according to the manufacturer’s protocol. Molecular identification of *An. gambiae* s.l was carried out using species-specific PCR with primers for *An. gambiae* s.s., *An. arabiensis* and *An. quadriannulatus* (19): AR-3T (5’-GTGTTAAGTGTCCTTCTCCGTC-3’; specific for *An. arabiensis*), GA-3T (5’-GCTTACTGGTTTGGTCGGCATGT-3; specific for *An. gambiae s*.*s*.), QD-3T (5’-GCATGTCCACCAACGTAAATCC-3’; specific for *An. quadriannulatus*) and IMP-UN (5’-GCTGCGAGTTGTAGAGATGCG-3’; common for all species). Each 25μl reaction volume contained 20-40ng of DNA, 5X Green GoTaq^®^ Reaction Buffer (Promega), 25mM MgCl_2_, 2mM of each dNTP, 1U GoTaq^®^ DNA polymerase and 25pmol/μl of primers AR-3T, GA-3T, QD-3T and IMP-UN. PCR cycling conditions were: 95°C for 5 minutes, followed by 30 amplification cycles (95°C for 30 seconds, 58°C for 30 seconds, 72°C for 30 seconds) and a final elongation step at 72°C for 5 minutes. Amplified PCR products were visualized on 1.5% agarose gels, stained with GelRed™ (Biotium, USA). Positive control DNA from *An. arabiensis* Sekoru, *An. gambiae* s.s. Kisumu and *An. quadriannulatus* Sangwe strains and no-template negative controls were included with all reaction runs. PCR products of 387bp, 463bp or 636bp were indicative of *An. arabiensis, An. gambiae* s.s. or *An. quadriannulatus*, respectively.

### Target site mutation detection

The presence of the G119S *Ace-1* mutation was determined using PCR restriction fragment length polymorphism analysis (59). Amplifications were performed in 25μl reactions containing 20-40ng of DNA, 5X Green GoTaq^®^ Reaction Buffer (Promega), 2.5mM of each dNTP, 1U GoTaq^®^ DNA polymerase, 25pmol/μl of primers MOUSTDIR1 (5’-CCGGGNGCSACYATGTGGAA-3’) and MOUSTREV1 (5’-ACGATMACGTTCTCYTCCGA-3’). PCR cycling conditions were 95°C for 5 minutes, followed by 35 amplification cycles (95°C for 30 seconds, 52°C for 30 seconds, 72°C for 1 minute) and a final elongation step at 72°C for 5 minutes. PCR products were initially visualized on 2% agarose gels, stained with GelRed™ (Biotium, USA) before incubation with *Alu*I restriction enzyme (New England Biolabs, USA) at 37°C for 16 hours, followed by 65°C for 20 minutes. DNA fragments were visualized on 2% agarose gels, stained with GelRed™ (Biotium, USA). DNA from *An. arabiensis* Sekoru was used as a negative control alongside a no-template control. DNA from *An. coluzzii* AKDR was used as a positive control. Undigested PCR products of 194bp indicated the susceptible allele (wild type) and 120bp and 74bp digested fragments indicated the presence of the resistant allele. The presence of all three bands indicated the sample was a heterozygote.

West African *kdr* (L1014S) and East African *kdr* (L1014F) alleles were detected using protocols for allele-specific PCR (AS-PCR) (60,61). Primers IPCF (5’-GATAATGTGGATAGATTCCCCGACCATG-3’), AltRev (5’-TGCCGTTGGTGCAGACAAGGATG -3’), WT-R (5’-GGTCCATGTTAATTTGCATTACTTACGAATA -3’) and East-F (5’-CTTGGCCACTGTAGTGATAGGAAAATC-3’) were used to detect the L1014S allele (AS-PCR East), whereas primers IPCF, AltRev, WT-R and West-F (5’-CTTGGCCACTGTAGTGATAGGAAATGTT-3’) were used to detect the L1014F allele (AS-PCR West). Each 25μl reaction volume contained 20-40ng of DNA, 5X Green GoTaq^®^ Reaction Buffer (Promega), 25mM MgCl_2_, 2mM of each dNTP, 1U GoTaq^®^ DNA polymerase, 2.5pmol/μl of primers IPCF and AltRev and either 5pmol/μl of primer WT-R and 2.5pmol/μl of primer East-F to detect the L1014S allele (AS-PCR East), or 25pmol/μl of primer WT-R and 8.8pmol/μl of primer West-F to detect the L1014F allele (AS-PCR West). PCR cycling conditions were 95°C for 5 minutes, followed by 35 amplification cycles (95°C for 30 seconds, 57°C for East or 59°C for West for 30 seconds, 72°C for 30 seconds) and a final elongation step at 72°C for 5 minutes. Amplified PCR products were visualized on 2% agarose gels, stained with GelRed™ (Biotium, USA). DNA from *An. gambiae* Kisumu was used as a negative control alongside a no-template control. DNA from *An. coluzzii* AKDR and *An. gambiae* s.s. RSP-ST were used as positive controls for L1014F and L1014S, respectively. Successful amplification was indicated by a PCR product of 314 bp; additional bands of 214bp and 156bp identified susceptible (wild type) and resistant alleles, respectively. Pearson’s Chi squared tests were used to evaluate deviations from Hardy-Weinberg equilibrium at the population-level.

### RNA extraction and cDNA library preparation

Total RNA was isolated from three pools containing five mosquitoes each from the following groups: mosquitoes phenotyped as resistant following a malathion or permethrin bioassay, non-insecticide exposed mosquitoes and susceptible *An. arabiensis* colony mosquitoes from Dongola and Sekoru strains. RNA was extracted using the Arcturus^®^ PicoPure^®^ RNA isolation kit (Life Technologies, USA) and quantified using the Agilent RNA ScreenTape 4200 assay, according to the manufacturers’ protocols. Two micrograms of starting material were treated with Baseline-ZERO™ DNase (Lucigen, USA) and ribosomal RNA was removed using the Ribo-Zero™ Magnetic Core Kit and Ribo-Zero™ rRNA Removal kit (Illumina, USA), according to the manufacturers’ protocols. Individual RNA-Seq libraries were prepared from each pool of extracted RNA using the ScriptSeq™ v2 RNA-Seq library preparation kit (Illumina, USA), using 12 cycles of PCR amplification, according to the manufacturer’s protocol. Libraries were purified using Agencourt AMPure XP beads (Beckman Coulter, USA) and assessed for quantity and size distribution using the Agilent DNA ScreenTape D5000 assay.

### RNA-sequencing, quality control and read mapping

Two experiments, each comprising nine RNA-Seq libraries, were sequenced as 2 × 125bp paired-end reads, on the Illumina HiSeq platform at the CDC. The first experiment (henceforth “malathion experiment”) contained three biological replicates each of malathion bioassay survivors, non-exposed mosquitoes and the susceptible Dongola strain. The second experiment (henceforth “permethrin experiment”) contained three biological replicates each of permethrin bioassay survivors, non-exposed mosquitoes and the susceptible Sekoru strain. Each experiment was sequenced on two HiSeq lanes to give an estimate of technical variation.

De-multiplexed paired end sequencing reads for each sample were evaluated for quality using FastQC v0.11.5 (62). Concatenated files for R1 and R2 reads were used for downstream analysis. Initially concatenated files for each sample were trimmed and filtered using fastp v0.21.0 (63) to remove adapter and low-quality reads according to the following criteria: minimum base quality score=20, minimum length required=25, polyG and poly tail trimming=True. Trimmed and filtered read pairs (R1/R2) were aligned against the reference genome, *An. arabiensis* Dongola (genome assembly version=AaraD1.11, GeneBank assembly identifier=GCA_000349185.1; GeneBank WGS Project=APCN01), directly downloaded from VectorBase (release 48) (64), using ‘subjunc’ v2.0.1, part of the subread aligner v2.0.1 (65), with default parameters. The resulting alignment was filtered to remove reads with low mapping quality (q< 10) and sorted successively using Samtools v1.10 (66). Descriptive statistics for the malathion and permethrin read libraries and sequencing alignments are shown in Table S1.

Tags (a read pair or single, unpaired read) mapped to the sense orientation of the annotated *An. arabiensis* Dongola genes (gene set of AaraD1.11 in gff downloaded from release 48 from Vector Base), were quantified using FeatureCounts, as part of the subread-aligner package v2.0.1 (65). The tag count with FeatureCount was carried out using the following criteria: 1) count only read pairs that have both ends aligned; 2) count fragment instead of reads; 3) minimum number of overlaps required=1; 4) feature_type=exon; 5) attribute type=gene_id; and 6) strandness=sense. The FeatureCount analysis generated a tag count matrix table which was inputted to edgeR (67) for differential expression analysis. Metrics describing the transcriptome alignments for the malathion and permethrin experiments are shown in Table S4.

### Differential transcription analysis and GO enrichment analysis

To remove the effect of noise and lowly expressed genes, for each pairwise comparison, genes with a total tag count less than 50 across all libraries (control *vs* treatment) were filtered out before further analysis. Only genes with a total tag count equal to or higher than 50 were considered. The function *calcNormFactors* (part of the edgeR package (67)), using the TMM (Trimmed Mean M-values) method, was used to normalize tag count among samples, by finding a set of scaling factors for the library sizes that minimized the log-fold changes between samples for most genes. The tag count was not normalized for gene length and GC content, as these values do not vary from sample to sample, so this would be expected to have little effect on DEGs. The DEGs between control (unexposed) and resistant (exposed) mosquitoes were selected after multiple testing using the *decideTests* function, part of the limma package (68). A critical value absolute fold-change=2 and FDR (False Discovery Rate) ≤ 0.01 was used. Different pairwise comparisons were conducted: 1) between resistant field mosquitoes (treatment) and unexposed field mosquitoes (control): CON-M *vs* MAL-R and CON-P *vs* PERM-R; 2) between a susceptible laboratory strain and exposed field mosquitoes: DON *vs* MAL-R, SEK *vs* PERM-R and DON *vs* PERM-R; 3) between the two susceptible laboratory strain: DON *vs* SEK; and 4) between field mosquitoes exposed to different insecticides: MAL-R vs PERM-R.

The annotation set of the AraD1.11 reference genome included 13,307 protein-coding genes and 378 additional non-coding genes (Table S8) (https://legacy.vectorbase.org/organisms/dongola/aarad111). However, Gene Ontology (GO) description of only 9,074 of these genes was provided in VectorBase (64) (cellular component: 4,784; molecular function: 7,261; biological processes: 5316). To increase the annotation efficiency, the predicted protein gene set fasta file of AraD1.11 was downloaded from VectorBase (release 48) (64) and was used for functional annotation using Blast2GO (69). A Blastp search of the protein fasta file was conducted against the Insecta category of the non-redundant protein NCBI database, with a maximum e-value cut-off of 1e−3. Additionally, the RefSeq protein IDs corresponding to the best blast hits of each query sequence were mapped to the GO database as curated and updated in the last release of Blast2GO database (November 2020). The resulting non-annotated genes from the Blast2GO analysis were mapped to the *An. gambiae* proteome (AgamP4.13) using a Blastp search with a maximum e-value cut-off of 1e−10 for ortholog inference. The best alignments (based on e-value and alignment score) were considered as orthologous genes, were ID mapped to the GO annotation of AgamP4.13 using the panda’s python library (70). The newly annotated genes were concatenated with the Blast2GO annotation, which was used as the background for the functional enrichment analysis of the DEGs. From this analysis, 10,456 (78.6%) of 13,307 protein coding genes were GO annotated.

GO term enrichment analysis of up- and down-regulated genes was carried out using Goatools (71) based on the go-basic database (release 2021-02-01). The list of 10,456 annotated genes of *An. arabiensis* with their associated GO terms was used as the background reference set. The *P* values used to evaluate significantly enriched GO terms were calculated based on Fisher’s exact test and corrected by Benjamini-Hochberg multiple test correction method. Finally, we used a FDR adjusted *P*-value <0.05 to tag statistically significant overrepresented GO terms associated with the list of DEGs.

### qRT-PCR validation of RNA-seq data

A subset of eleven differentially transcribed genes was selected for quantitative real-time reverse transcription PCR validation (qRT-PCR). One microgram of RNA from three replicates of malathion resistant or permethrin resistant, non-exposed and Dongola strain mosquitoes were used to synthesize cDNA using the High-Capacity cDNA Reverse Transcription kit (Applied Biosystems, USA) with oligo-dT20 (New England Biolabs, USA), according to the manufacturer’s instructions. Primer sequences and efficiencies are detailed in Table S9. Standard curves of Ct values for each gene were generated using a five-fold serial dilution of cDNA to assess PCR efficiency. Reactions were performed using either a QuantStudio 6 Flex Real-Time PCR system (Applied Biosystems, USA) with PowerUp SYBR Green Master Mix (Applied Biosystems, USA) or a Stratagene Mx3005P Real-Time PCR system (Agilent Technologies) with LightCycler^®^ 480 SYBR Green I Master Mix (Roche, UK). cDNA from each sample was used as a template in a three-step reaction: 50°C for 2 minutes, denaturation at 95°C for 10 minutes, followed by 40 cycles of 15 seconds at 95°C, 1 minute at 60°C and a final step of 15 seconds at 95°C, 1 minute at 60°C, and 15 seconds at 95°C. The relative expression level and Fold Change (FC) of each target gene from resistant field samples, relative to the susceptible laboratory strain (Dongola), were calculated using the 2^-ΔΔCT^ method (72), incorporating PCR efficiency. Two housekeeping genes, ribosomal protein S7 (RpS7: AARA000046) and ubiquitin (AARA016296), were used for normalisation.

### Sequence polymorphism analysis

The RNA-Seq reads of all resistant groups and susceptible strains were mined for the prevalence of non-synonymous Single Nucleotide Polymorphisms (SNPs) involved in *Anopheles spp* resistance to either DDT, organophosphate or pyrethroid insecticides. The primary target of the analysis was the *para* Voltage-Gated Sodium Channel (*VGSC*) gene (AARA017729), for which the presence of 21 recently reported non-synonymous SNPs (A1125V, A1746S, A1934V, D466H, E1597G, F1920S, I1527T, I1868T, I1940T, K1603T, L995F, L995S, M490I, N1575Y, P1874L, P1874S, T791M, V1254I, V1853I, V402L, and V1853I) were investigated (28). Additionally, non-synonymous variants G119S in the acetylcholinesterase (*Ace-1*) gene (AARA001814), L119F and I114T in GSTe2 (AARA008732) (23,24), were also investigated. Prevalence of the target site mutations in the RNA-Seq datasets was determined as follows. The coding sequences (CDS) corresponding to VGSC, Ace-1, and GSTe2 from AaraD1.11 were downloaded from VectorBase (64) and were aligned separately with their respective homologous gene retrieved from the AgamP4.4 gene set, using Clustalw Omega (73). Next, the sequence (∼30-40 nucleotides) flanking the codon and the site of interest from each gene in *An. arabiensis* was identified and extracted from the alignment as described here (74). The resulting flanking sequence was BLASTn (75) searched against the AaraD1.11 reference genome (release 48 in Vectorbase) (64), which gave the exact chromosomal numerical position of the nucleotide. Finally, the sorted bam files, which were previously used as the input featureCount for DEG analysis were separately uploaded to Integrative Genomics Viewer (IGV) (76) and zoomed to the position to the flanking sequence. The allele frequency in the population was calculated as the percentage of RNA-Seq reads spanning the codon with the SNP of interest.

## Declarations

### Ethics approval and consent to participate

The study protocol was reviewed and approved by the institutional review boards (IRBs) of the Institute of Health, Jimma University (THRPGD/843/17) and the U.S. Centers for Disease Control and Prevention, USA (2017-227).

### Consent for publication

Not applicable

### Availability of data and material

Sequence data generated by this study is available at Sequence Read Archive (SRA) BioProject PRJNA730212 (accession numbers: SAMN19223816-SAMN19223833). All other relevant data are available from the corresponding author upon reasonable request.

### Author contributions

LAM, LMI, DY, SI and AL designed the study. LAM and DY conducted the field work. LAM and LMI undertook mosquito rearing, phenotyping and preparation of samples for sequencing. Laboratory supervision was provided by AL. DD, LAM and LMI performed the formal data analysis. LAM, LMI, DD and AL drafted the manuscript, which was reviewed by DY and SI.

## Acknowledgements

The authors would like to thank all of the entomology fieldworkers of the Tropical and Infectious Diseases Research Center (TIDRC), Jimma University for their dedicated work and the residents of Asendabo for their study participation. We gratefully acknowledge members of the Biotechnology Core Facility Branch, U.S. Centers for Disease Control and Prevention (CDC), Atlanta. We thank Dustin Miller for providing PCR controls, Gareth Weedall and Steven E. Massey for analytical expertise and Yikun Li for PCR technical advice.

## Disclaimer

The findings and conclusions in this paper are those of the authors and do not necessarily represent the official position of the Centers for Disease Control and Prevention.

## Supplementary information

**Table S1**. RNA sequencing and mapping statistics for the malathion and permethrin experiments.

**Table S2**. Full results of pairwise differential gene expression analyses of *Anopheles arabiensis* RNA-seq datasets.

**Table S3**. Full results of gene ontology enrichment analyses for differentially expressed gene sets.

**Table S4**. Metrics describing the transcriptome alignments for the malathion and permethrin experiments.

**Table S5**. Full results of core DEGs shared between MAL-R *vs* DON and PERM-R *vs* DON conditions.

**Table S6**. Target site mutation population allele frequencies.

**Table S7**. Target site mutation allele counts.

**Table S8**. Functional annotation of AraD1.11 protein coding genes.

**Table S9**. Oligonucleotide primers used for qRT-PCR.

**Figure S1**. Volcano plots of gene expression for different pairwise comparisons between conditions. Detoxification gene families are indicated in red (COE: carboxylesterases), blue (CYP: cytochrome P450s) and black (GST: glutathione-S-transferases). Cuticular proteins are indicated in green (CP) and salivary gland proteins are indicated in pink (SP). In each plot, genes overexpressed in the population are >0 on the x-axis. Vertical dotted lines indicate 2-fold expression differences and the horizontal line indicates a *P*-value of 0.01.

**Figure S2**. Experimental design and differentially expressed transcripts among resistant (R), susceptible (S) and unexposed (C) mosquito groups in the permethrin experiment relative to the SEK susceptible strain. Each Venn diagram section shows the number of differentially expressed genes meeting each set of conditions (*P*-values were adjusted for multiple testing based on FDR<0.01 and FC>2). For a list of all DEGs for each comparison see Table S2.

**Figure S3**. Multidimensional scaling plot of all mosquito groups.

## References

1. World Health Organization. World Malaria Report 2020. Geneva;2020.

2. Presidents Malaria Initiative. President’s Malaria Initiative Ethiopia Malaria Operational Plan FY 2019. CDC;2019.

3. Messenger LA, Shililu J, Irish SR, Anshebo GY, Tesfaye AG, Ye-Ebiyo Y, Chibsa S, Dengela D, Dissanayake G, Kebede E, Zemene E, Asale A, Yohannes M, Taffese HS, George K, Fornadel C, Seyoum A, Writz RA, Yewhalaw D. 2017. Insecticide resistance in Anopheles arabiensis from Ethiopia (2012–2016): a nationwide study for insecticide resistance monitoring. Malar J 16:469. doi:10.1186/s12936-017-2115-2.

4. Alemayehu E, Asale A, Eba K, Getahun K, Tushune K, Bryon A, Morou E, Vontas J, Van Leeuwen T, Duchateau L, Yewhalaw D. 2017. Mapping insecticide resistance and characterization of resistance mechanisms in Anopheles arabiensis (Diptera: Culicidae) in Ethiopia. Parasit Vectors 10(1):407. doi:10.1186/s13071-017-2342-y.

5. Yewhalaw D, Van Bortel W, Denis L, Coosemans M, Duchateau L, Speybroeck N. 2010. First evidence of high knockdown resistance frequency in Anopheles arabiensis (Diptera: Culicidae) from Ethiopia. Am J Trop Med Hyg 83(1):122–5. doi: 10.4269/ajtmh.2010.09-0738.

6. Birhanu A Asale A, Yewhalaw D. 2019. Bio-efficacy and physical integrity of piperonylbutoxide coated combination net (PermaNet® 3.0) against pyrethroid resistant population of Anopheles gambiae s.l. and Culex quinquefasciatus mosquitoes in Ethiopia. Malar J 18:224. doi: 10.1186/s12936-019-2641-1.

7. Müller P, Warr E, Stevenson BJ, Pignatelli PM, Morgan JC, Steven A, Yawson AE, Mitchell SN, Ranson H, Hemingway J, Paine MJI, Donnelly MJ. 2008. Field-caught permethrin-resistant Anopheles gambiae overexpress CYP6P3, a P450 that metabolises pyrethroids. PLoS Genet 4(11):e1000286. doi: 10.1371/journal.pgen.1000286.

8. Stevenson BJ, Bibby J, Pignatelli P, Muangnoicharoen S, O’Neill PM, Lian LY, Müller P, Nikou D, Steven A, Hemingway J, Sutcliffe MJ, Paine MJI. 2011. Cytochrome P450 6M2 from the malaria vector Anopheles gambiae metabolizes pyrethroids: Sequential metabolism of deltamethrin revealed. Insect Biochem Mol Biol 41(7):492–502.

9. Chiu T-L, Wen Z, Rupasinghe SG, Schuler MA. 2008. Comparative molecular modeling of Anopheles gambiae CYP6Z1, a mosquito P450 capable of metabolizing DDT. Proc Natl Acad Sci U S A105(26):8855–60. doi: 10.1073/pnas.0709249105.

10. Ibrahim SS, Riveron JM, Stott R, Irving H, Wondji CS. 2016. The cytochrome P450 CYP6P4 is responsible for the high pyrethroid resistance in knockdown resistance-free Anopheles arabiensis. Insect Biochem Mol Biol 68:23–32. doi:10.1016/j.ibmb.2015.10.015.

11. Riveron JM, Yunta C, Ibrahim SS, Djouaka R, Irving H, Menze BD, Ismail HM, Hemingway J, Ranson H, Albert A, Wondji CS. 2014. A single mutation in the GSTe2 gene allows tracking of metabolically based insecticide resistance in a major malaria vector. Genome Biol 15(2):R27. doi:10.1186/gb-2014-15-2-r27.

12. Ingham VA, Wagstaff S, Ranson H. 2018. Transcriptomic meta-signatures identified in Anopheles gambiae populations reveal previously undetected insecticide resistance mechanisms. Nat Commun 9:5282. doi: 10.1038/s41467-018-07615-x.

13. Ingham VA, Pignatelli P, Moore JD, Wagstaff S, Ranson H. 2017. The transcription factor Maf-S regulates metabolic resistance to insecticides in the malaria vector Anopheles gambiae. BMC Genomics 18(1):669. doi:10.1186/s12864-017-4086-7.

14. Isaacs AT, Mawejje HD, Tomlinson S, Rigden DJ, Donnelly MJ. 2018. Genome-wide transcriptional analyses in Anopheles mosquitoes reveal an unexpected association between salivary gland gene expression and insecticide resistance. BMC Genomics 19(1):225. doi: 10.1186/s12864-018-4605-1.

15. Ingham VA, Anthousi A, Douris V, Harding NJ, Lycett F, Morris M, Vontas J, Ranson H. 2019. A sensory appendage protein protects malaria vectors from pyrethroids. Nature 577:376–80. doi: 10.1038/s41586-019-1864-1.

16. Balabanidou V, Kampouraki A, MacLean M, Blomquist GJ, Tittiger C, Juárez MP, Mijailovsky SJ, Chalepakis G, Anthousi A, Lynd A, Antoine S, Hemingway J, Ranson H, Lycett GJ, Vontas J. 2016. Cytochrome P450 associated with insecticide resistance catalyzes cuticular hydrocarbon production in Anopheles gambiae. Proc Natl Acad Sci U S A 113(33):9268–73. doi: 10.1073/pnas.1608295113.

17. Simma EA, Dermauw W, Balabanidou V, Snoeck S, Bryon A, Clark RM, Yewhalaw D, Vontas J, Duchateau L, Van Leeuwen T. 2019. Genome-wide gene expression profiling reveals that cuticle alterations and P450 detoxification are associated with pyrethroid resistance in Anopheles arabiensis populations from Ethiopia. Pest Manag Sci 75(7):1808–18. doi: 10.1003/ps.5374.

18. Morgan JC, Irving H, Okedi LM, Steven A, Wondji CS. 2010. Pyrethroid resistance in an Anopheles funestus population from Uganda. PLoS One 5(7):1–8. doi: 10.1371/journal/pone.0011872.

19. Wilkins EE, Howell PI, Benedict MQ. 2006. IMP PCR primers detect single nucleotide polymorphisms for Anopheles gambiae species identification, Mopti and Savanna rDNA types, and resistance to dieldrin in Anopheles arabiensis. Malar J 5:125. doi: 10.1186/1475-2875-5-125.

20. Centers for Disease Control and Prevention. Guideline for Evaluating Insecticide Resistance in Vectors Using the CDC Bottle Bioassay. CDC Methods. 2012;1–28.

21. Adedeji EO, Ogunlana OO, Fatumo S, Beder T, Ajamma Y, Koenig R, Adebiyi E. 2020. Anopheles metabolic proteins in malaria transmission, prevention and control: a review. Parasit Vectors 13:465. doi: 10.1186/s13071-020-04342-5.

22. Robinson MD, McCarthy DJ, Smyth GK. 2010. edgeR: a Bioconductor package for differential expression analysis of digital gene expression data. Bioinformatics 26(1):139–40. doi: 10.1093/bioinformatics/btp616.

23. Mitchell SN, Rigden DJ, Dowd AJ, Lu F, Wilding CS, Weetman D, Dadzie S, Jenkins AM, Regna K, Boko P, Djogbenou L, Muskavitch MAT, Ranson H, Paine MJI, Mayans O, Donnelly MJ. 2014. Metabolic and target-site mechanisms combine to confer strong DDT resistance in Anopheles gambiae. PLoS One 9(3):e92662. doi: 10.1371/journal.pone.0092662.

24. Lucas ER, Rockett KA, Lynd A, Essandoh J, Grisales N, Kemei B, Njoroge H, Hubbart C, Rippon EJ, Morgan J, Van’t Hof AE, Ochomo E, Kwiatkowski DP, Weetman D, Donnelly MJ. 2019. A high throughput multi-locus insecticide resistance marker panel for tracking resistance emergence and spread in Anopheles gambiae. Sci Rep 9:13335. doi: 10.1038/s41598-019-49892-6.

25. Jones CM, Liyanapathirana M, Agossa FR, Weetman D, Ranson H, Donnelly MJ, Wilding CS. 2012. Footprints of positive selection associated with a mutation (N1575Y) in the voltage-gated sodium channel of Anopheles gambiae. Proc Natl Acad Sci U S A 109(17):6614–9.

26. Collins E, Vaselli NM, Sylla M, Beavogui AH, Orsborne J, Lawrence G, Wiegand RE, Irish SR, Walker T, Messenger LA. 2019. The relationship between insecticide resistance, mosquito age and malaria prevalence in Anopheles gambiae s.l. from Guinea. Sci Rep 9(1):8846.

27. Lynd A, Oruni A, Van’t Hof A, Morgan JC, Naego LB, Pipini D, O’Kines KA, Bobanga TL, Donnelly MJ, Weetman D. 2018. Insecticide resistance in Anopheles gambiae from the northern Democratic Republic of Congo, with extreme knockdown resistance (kdr) mutation frequencies revealed by a new diagnostic assay. Malar J 17(1):412.

28. Clarkson CS, Miles A, Harding NJ, O’Reilly AO, Weetman D, Kwiatkowski D, Donnelly MJ, Anopheles gambiae 1000 Genomes Consortium. 2021. The genetic architecture of target-site resistance to pyrethroid insecticides in the African malaria vectors Anopheles gambiae and Anopheles coluzzii. Mol Ecol doi: 10.1111/mec.15845.

29. Dagg K, Irish S, Wiegand RE, Shililu J, Yewhalaw D, Messenger LA. 2019. Evalution of toxicity of clothianidin (neonicotinoid) and chlorfenapyr (pyrrole) insecticides and cross-resistance to other public health insecticides in Anophles arabiensis from Ethiopia. Malar J 18:49. doi: 10.1186/s12936-019-2685-2.

30. Wondji CS, Coleman M, Kleinschmidt I, Mzilahowa T, Irving H, Ndula M, Rehman A, Morgan J, Barnes KG, Hemingway J. 2012. Impact of pyrethroid resistance on operational malaria control in Malawi. Proc Soc Natl Acad Sci U S A 109(47):19063–19070.

31. Abdalla H, Wilding CS, Nardini L, Pignatelli P, Koekemoer LL, Ranson H, Coetzee M. 2014. Insecticide resistance in Anopheles arabiensis in Sudan: temporal trends and underlying mechanisms. Parasit Vectors 7:213. doi: 10.1186/1756-3305-7-213.

32. Mitchell SN, Stevenson BJ, Müller P, Wilding CS, Egyir-Yawson A, Field SG, Hemingway J, Paine MJI, Ranson H, Donnelly MJ. 2012. Identification and validation of a gene causing cross-resistance between insecticide classes in Anopheles gambiae from Ghana. Proc Natl Acad Sci U S A 109(16):6147–52. doi: 10.1073/pnas.1203452109.

33. Vontas J, Grigoraki L, Morgan J, Tsakireli D, Fuseini G, Segura L, de Carvalho JN, Nguema R, Weetman D, Slotman MA, Hemingway J. 2018. Rapid selection of a pyrethroid metabolic enzyme CYP9K1 by operational malaria control activities. Proc Natl Acad Sci U S A 115(18):4619–4624. doi: 10.1073/pnas.1719663115.

34. Yunta C, Hemmings K, Stevenson B, Koekemoer LL, Matambo T, Pignatelli P, Voice M, Nasz S, Paine MJI. 2019. Cross-resistance profiles of malaria mosquitp P450s associated with pyrethroid resistance against WHO insecticides. Pestic Biochem Physiol 161:61–7. doi: 10.1016/j.pestbp.2019.06.007.

35. Vontas J, Katsavou E, Mavridis K. 2020. Cytochrome P450-based metabolic insecticide resistance in Anopheles and Aedes mosquito vectors: muddying the waters. Pestic Biochem Physiol 170:104666. doi: 10.1186/s12864-021-07646-7.

36. Voice MW, Kaaz AW, Peet CF, Paine MJ. 2012. Recombinant CYP6M2 inhibition by insecticides recommended by WHO for indoor residual spraying. Drug Metab Rev; doi: 10.3109/03602532.2012.744573.

37. Adolfi A, Poulton B, Anthousi A, Macilwee S, Ranson H, Lycett GJ. 2019. Functional genetic validation of key genes conferring insecticide resisatance in the major African malaria vector, Anopheles gambiae. Proc Natl Acad Sci U S A 116(51):25764–25772. doi: 10.1073/pnas.1914633116.

38. Bonizzoni M, Ochomo E, Dunn WA, Britton M, Afrane Y, Zhou G, Hartsel J, Lee MC, Xu J, Githeko A, Fass J, Yan G. 2015. RNA-seq analyses of changes in the Anopheles gambiae transcriptome associated with resistance to pyrethroids in Kenya: identification of candidate-resistance genes and candidate-resistance SNPs. Parasit Vectors 8:474. doi: 10.1186/s13071-015-1083-z.

39. Müller P, Chouaïbou M, Pignatelli P, Etang J, Walker ED, Donnelly MJ, Simard F, Ranson H. 2008. Pyrethroid tolerance is associated with elevated expression of antioxidants and agricultural practice in Anopheles arabiensis sampled from an area of cotton fields in Northern Cameroon. Mol Ecol 17(4):1145–55. doi: 10.1111/j.1365-294X.2007.03617.x.

40. Jones CM, Haji KA, Khatib BO, Bagi J, Mcha J, Devine GJ, Daley M, Kabula B, Ali AS, Majambere S, Ranson H. 2013. The dynamics of pyrethroid resistance in Anopheles arabiensis from Zanzibar and an assessment of the underlying genetic basis. Parasit Vectors 6:343. doi: 10.1186/1756-3305-6-343.

41. Abose T, Yeebiyo Y, Olana D, Alamirew D, Beyene Y, Regassa L. Reorientation and definition of the role of malaria vector control in Ethiopia: the epidemiology and control of malaria with special emphasis on the distribution, behaviour and susceptibility of insecticides of anopheline vectors and chloroquine resistance in Zwai, Central Ethiopia and other areas. WHO/MAL/98.1085;1998.

42. Yewhalaw D, Wassie F, Steurbaut W, Spanoghe P, Van Bortel W, Denis L, Tessema DA, Getachew Y, Coosemans M, Duchateau L, Speybroeck N. Multiple insecticide resistance: an impediment to insecticide-based malaria vector control program. PLoS One. 2011;6(1):e16066. doi: 10.1371/journal.pone.0016066.

43. Mekuriaw W, Yewhalaw D, Woyessa A, Dugassa S, Taffese H, Bashaye S, Nigatu W, Massebo F. 2019. Distribution and trends of insecticide resistance in malaria vectors in Ethiopia (1986-2017): a review. Ethiopian Journal of Public Health and Nutrition. 3:51–61.

44. Grossman MK, Uc-Puc V, Rodriguez J, Cutler DJ, Morran LT, Manrique-Saide P, Vazquez-Prokopec GM. 2018. Restoration of pyrethroid susceptibility in a highly resistant Aedes aegypti population. Biol Lett 14(6):20180022. doi: 10.1098/rsbl.2018.0022.

45. Shi L, Hu H, Zhou D, Yu J, Zhong D, Fang F, Chang X, Hu S, Zou F, Wang W, Sun Y, Shen B, Zhang D, Ma L, Zhou G, Yan G, Zhu C. 2015. Development of resistance to pyrethroid in Culex pipiens pallens populations under different insecticide selection pressures. PLoS Negl Trop Dis 9(8):e0003928. doi: 10.1371/journal.pntd.0003928.

46. Williams J, Flood L, Praulins G, Ingham VA, Morgan J, Lees RS, Ranson H. 2019. Characterisation of Anopheles strains used for laboratory screening of new vector control products. Parasit Vectors 12(1):522. doi.10.1186/s13071-019-3774-3.

47. Grau-Bové X, Tomlinson S, O’Reilly AO, Harding NJ, Miles A, Kwiatkowski D, Donnelly MJ, Weetman D and The Anopheles gambiae 1000 Genomes Consortium. 2020. Evolution of the insecticide target rdl in African Anopheles is driven by interspecific and interkaryotypic introgression. Mol Biol Evol 37(10):2900–917. doi.10.1093/molbev/msaa128.

48. Machani MG, Ochomo E, Zhong D, Zhou G, Wang X, Githeko AK, Yan G, Afrane YA. 2020. Phenotypic, genotypic and biochemical changes during pyrethroid resistance selection in Anopheles gambiae mosquitoes. Sci Rep 10:19063. doi: 10.1038/s41598-020-75865-1.

49. Balabanidou V, Kefi M, Aivaliotis M, Koidou V, Girotti JR, Mijalovsky SJ, Juarez MP, Papadogiorgaki E, Chalepakis G, Kampouraki A, Nikolaou C, Ranson H, Vontas J. 2019. Mosquitoes cloak their legs to resist insecticides. Proc R Soc B 286:20191091. doi: 10.1098/rspb.2019/1091.

50. Balabanidou V, Kampouraki A, MacLean M, Blomquist GJ, Tittiger C, Juarez MP, Mijailovsky SJ, Chalepakis G, Anthousi A, Lynd A, Antoine S, Hemingway J, Ranson H, Lycett GJ, Vontas J. 2016. Cytochrome P450 associated with insecticide resistance catalyzes cuticular hydrocarbon production in Anopheles gambiae. Proc Natl Acad Sci U S A 113(33):9268–9273. doi: 10.1073/pnas.1608295113.

51. Yahouedo GA, Chandre F, Rossignol M, Ginibre C, Balabanidou V, Mendez NGA, Pigeon O, Vontas J, Cornelie S. 2017. Contribution of cuticular permeability and enzyme detoxification to pyrethroid resistance in the major malaria vector Anopheles gambiae. Sci Rep 7:11091. doi: 10.1038/s41598-017-11357-z.

52. Mavridis K, Wipf N, Medves S, Erquiaga I, Müller P, Vontas J. 2019. Rapid multiplex gene expression assays for monitoring metabolic resistance in the major malaria vector Anopheles gambiae. Parasit Vectors 12:9. doi: 10.1186/s13071-018-3253-2.

53. Merzendorfer H, Zimoch L. 2003. Chitin metabolism in insects: structure, function and regulation of chitin synthases and chitinases. J Exp Biol 206(Pt 24):4393–412. doi: 10.1242/jeb.00709.

54. Wilding CS, Weetman D, Rippon EJ, et al. 2015. Parallel evolution or purifying selection, not introgression, explains similarity in the pyrethroid detoxification linked GSTE4 of Anopheles gambiae and An. arabiensis. Mol Genet Genomics 290(1):201–15. doi: 10.1007/s00438-014-0910-9.

55. Ibrahim SS, Ndula M, Riveron JM, et al. 2016. The P450 CYP6Z1 confers carbamate/pyrethroid cross-resistance in a major African malaria vector besides a novel carbamate-insensitive N485I acetylcholinesterase-1 mutation. Mol Ecol 25(14):3436–52. doi: 10.1111/mec.13673.

56. Gillies MT, Coetzee M. 1987. A supplement of the Anophelinae of Africa south of the Sahara (Afrotropical Region). Publ South African Inst Med Res No 55.

57. Balkew M, Ibrahim M, Koekemoer LL, Brooke BD, Engers H, Aseffa A, Gebre-Michael T, Elhassen I. 2010. Insecticide resistance in Anopheles arabiensis (Diptera: Culicidae) from villages in central, northern and south west Ethiopia and detection of kdr mutation. Parsit Vectors 3:40. doi: 10.1186/1756-3305-3-40.

58. World Health Organization. 2013. Test procedures for insecticide resistance monitoring in malaria vector mosquitoes. World Heal Organ Tech Rep Ser 22.

59. Weill M, Malcolm C, Chandre F, Mogensen K, Berthomieu A, Marquine M, Raymond M. 2004. The unique mutation in ace-1 giving high insecticide resistance is easily detectable in mosquito vectors. Insect Mol Biol 13(1):1–7. doi: 10.1111/j.1365-2583.2004.00452.x.

60. Martinez-Torres D, Chandre F, Williamson MS, Darriet F, Bergé JB, Devonshire AL, Guillet P, Pasteur N PD. 1998. Molecular characterization of pyrethroid knockdown resistance (kdr) in the major malaria vector Anopheles gambiae s.s. Insect Mol Biol 7:179–84. doi: 10.1046/j.1365-2583.1998.72062.x.

61. Ranson H, Jensen B, Vulule JM, Wang X, Hemingway J, Collins FH. 2000. Identification of a point mutation in the voltage-gated sodium channel gene of Kenyan Anopheles gambiae associated with resistance to DDT and pyrethroids. Insect Mol Biol 9(5):491–7. doi: 10.1046/j.1365-2583.2000.00209.x.

62. Andrews S. 2016. FastQC: A quality control tool for high throughput sequence data. Babraham Bioinforma.

63. Chen S, Zhou Y, Chen Y, Gu J. 2018. fastp: an ultra-fast-all-in-one FASTQ preprocessor. Bioinformatics 34(17):i884–i890. soi: 10.1093/bioinformatics/bty560.

64. Giraldo-Calderón GI, Emrich SJ, MacCallum RM, Maslen G, Dialynas E, Topalis P, Ho N, Gesing S, VectorBase Consortium; Madey G, Collins FH, Lawson D. 2015. VectorBase: An updated Bioinformatics Resource for invertebrate vectors and other organisms related with human diseases. Nucleic Acids Res 43:D707–13. doi: 10.1093/nar/gku1117.

65. Liao Y, Smyth GK, Shi W. 2013 The Subread aligner: fast, accurate and scalable read mapping by seed-and-vote. Nucleic Acids Res 41(10):e108. doi: 10.1093/nar/gkt214.

66. Li H, Handsaker B, Wysoker A, Fennell T, Ruan J, Homer N, Marth G, Abecasis G, Durbin R and 1000 Genome Project Data Processing Subgroup. 2009. The Sequence Alignment/Map format and SAMtools. Bioinformatics 25(16):2078–2079. doi: 10.1093/bioinformatics/btp352.

67. Robinson MD, McCarthy DJ, Smyth GK. 2010. edgeR: a Bioconductor package for differential expression analysis of digital gene expression data. Bioinformatics 26(1):139–140. doi: 10.1093/bioinformatics/btp616.

68. Ritchie ME, Phipson B, Wu D, Hu Y, Law CW, Shi W, Smyth GK. 2015. Limma powers differential expression analyses for RNA-sequencing and microarray studies. Nucleic Acids Res 43(7):e47. doi: 10.1093/nar/gkv007.

69. Conesa A, Götz S. 2008. Blast2GO: a comprehensive suite for functional analysis in plant genomics. Int J Plant Genomics 2008:619832. doi: 10.1155/2008/619832.

70. McKinney W. 2011. pandas: a foundational Python library for data analysis and statistics.

71. Klopfenstein DV, Zhang L, Pedersen BS, Ramirez F, Warwick Vesztrocy A, Naldi A, Mungall CJ, Yunes JM, Botvinnik O, Weigel M, Dampier W, Dessimoz C, Flick P, Tang H. 2018. GOATOOLS: a Python library for Gene Ontology. Sci Rep 8:10872. doi: 10.1038/s41598-018-28948-z.

72. Rao X, Huang X, Zhou Z, Lin X. 2013. An improvement of the 2^(–delta delta CT) method for quantitative real-time polymerase chain reaction data analysis. Biostat Bioinforma Biomath 3:71–85.

73. Sievers F, Wilm A, Dineen D, Gibson TJ, Karplus K, Li W, Lopez R, McWilliam H, Remmert M, Söding J, Thompson JD, Higgins DG. 2011. Fast, scalable generation of high-quality protein multiple sequence alignments using Clustal Omega. Mol Syst Biol 7:539. doi: 10.1038/msb.2011.75

74. Lol JC, Castañeda D, Mackenzie-Impoinvil L, Romero CG, Lenhart A, Padilla NR. 2019. Development of molecular assays to detect target-site mechanisms associated with insecticide resistance in malaria vectors from Latin America. Malar J 18:202. doi: 10.1186/s12936-019-2834-7.

75. Chen Y, Ye W, Zhang Y, Xu Y. 2015. High speed BLASTN: an accelerated MegaBLAST search tool. Nucleic Acids Res 43(16):7762–8. doi: 10.1093/nar/gkv784.

76. Thorvaldsdótti H, Robinson JT, Mesirov JP. 2013. Integrative Genomics Viewer (IGV): high-performance genomics data visualization and exploration. Brief Bioinform 14(2):178–92. doi: 10.1093/bib/bbs017.

